# Bivariate genomic prediction of phenotypes by selecting epistatic interactions across years

**DOI:** 10.1101/2020.11.18.388330

**Authors:** Elaheh Vojgani, Torsten Pook, Armin C. Hölker, Manfred Mayer, Chris-Carolin Schön, Henner Simianer

## Abstract

The importance of accurate genomic prediction of phenotypes in plant breeding is undeniable, as higher prediction accuracy can increase selection responses. In this study, we investigated the ability of three models to improve prediction accuracy by including phenotypic information from the last growing season. This was done by considering a single biological trait in two growing seasons (2017 and 2018) as separate traits in a multi-trait model. Thus, bivariate variants of the Genomic Best Linear Unbiased Prediction (GBLUP) as an additive model, Epistatic Random Regression BLUP (ERRBLUP) and selective Epistatic Random Regression BLUP (sERRBLUP) as epistasis models were compared with respect to their prediction accuracies for the second year. The results indicate that bivariate ERRBLUP is slightly superior to bivariate GBLUP in predication accuracy, while bivariate sERRBLUP has the highest prediction accuracy in most cases. The average relative increase in prediction accuracy from bivariate GBLUP to maximum bivariate sERRBLUP across eight phenotypic traits and studied dataset from 471/402 doubled haploid lines in the European maize landrace Kemater Landmais Gelb/Petkuser Ferdinand Rot, were 7.61 and 3.47 percent, respectively. We further investigated the genomic correlation, phenotypic correlation and trait heritability as the factors affecting the bivariate model’s predication accuracy, with genetic correlation between growing seasons being the most important one. For all three considered model architectures results were far worse when using a univariate version of the model, e.g. with an average reduction in prediction accuracy of 0.23/0.14 for Kemater/Petkuser when using univariate GBLUP.

**Key Massage:** Bivariate models based on selected subsets of pairwise SNP interactions can increase the prediction accuracy by utilizing phenotypic data across years under the assumption of high genomic correlation across years.

## Introduction

In plant breeding, genomic prediction has become a daily tool (Bernal-Vasquez *et al.* 2014; Stich and Ingheland 2018) which enables the optimization of phenotyping costs of breeding programs (Akdemir and Isidro-Sánchez 2019). The importance of genomic prediction of phenotypes is not restricted to plants. Livestock (Daetwyler *et al.* 2013) and human research (de los Campos *et al.* 2013) also have been widely developed in this regard. In the context of plant and animal breeding, accurately predicting phenotypic traits is of special importance, since raising all animals and growing all crops to measure their performances requires a considerable amount of money under limited resources (Martini *et al.* 2016).

Several statistical models have been compared over the last decades in the term of prediction accuracy. In this context, genomic best linear unbiased prediction (GBLUP) (Meuwissen *et al.* 2001; VanRaden 2007) as an additive linear mixed model has been widely used due to its high robustness, computing speed and superiority in predictive ability to alternative prediction models like Bayesian methods, especially in small reference populations (Da *et al.* 2014; Rönnegård and Shen 2016; Covarrubias-Pazaran *et al.* 2018; Wang *et al.* 2018). Furthermore, inclusion of genotype ×environment interaction into additive genomic prediction models can result in an increase in prediction accuracy (Hallauer *et al.* 2010; Bajgain *et al.* 2020). Such approaches allow borrowing information across environments which potentially leads to higher accuracy in phenotype prediction in multi environment models (Burgueño *et al.* 2012). In fact, multivariate mixed models have been originally proposed in the context of animal breeding (Henderson and Quaas 1976) with the purpose of modeling the genomic correlation among traits, longitudinal data, and modeling genotype by environment interactions across multiple years or environments (Mrode 2014; Lee and van der Werf 2016; Covarrubias-Pazaran *et al.* 2018). A multivariate GBLUP model was reported to have higher prediction accuracy than univariate GBLUP (Jia and Jannink 2012) when the genetic correlations were medium (0.6) or high (0.9) (Covarrubias-Pazaran *et al.* 2018). It was also shown that aggregating the phenotypic data over years to train the model and predict the performance of lines in the following years is a possible approach which can improve prediction accuracy (Auinger *et al.* 2016; Schrag *et al.* 2019a).

In addition, inclusion of epistasis, defined as the interaction between loci (Falconer and Mackay 1996; Lynch and Walsh 1998), into the genomic prediction model results in more accurate phenotype prediction (Hu *et al.* 2011; Wang *et al.* 2012; Mackay 2014; Martini *et al.* 2016; Vojgani *et al.* 2019b) due to the considerable contribution of epistasis in genetic variation of quantitative traits (Mackay 2014). In this context, several statistical models have been proposed. Extended genomic best linear unbiased prediction (EG-BLUP, Jiang and Reif 2015) and categorical epistasis (CE, Martini *et al.* 2017) models are using a marker-based epistatic relationship matrix that is constructed in a highly efficient manner. It has been shown that the CE model is as good as or better than EG-BLUP and does not possess undesirable features of EG-BLUP such as coding-dependency (Martini *et al.* 2017).

Moreover, it was shown that the accuracy of the epistasis genomic prediction model can be increased in one environment by variable selection in another environment (Martini *et al.* 2016). In this approach, the full epistasis model was reduced to a model with a subset of the largest epistatic interaction effects, resulting in an increase in predictive ability (Martini *et al.* 2016), through borrowing information across environments. Vojgani *et al.* (2019b) showed that the prediction accuracy can be increased even further by selecting the interactions with the highest absolute effect sizes / variances in the epistasis model. Resulting higher computational needs were offset by the development of a highly efficient software package (Vojgani *et al.* 2019a) to perform computations in a bit-wise manner (Schlather 2020). Thus, enabling to conduct such predictions with data sets of practically relevant size across environments in the same year, both with respect to sample size and number of markers (Vojgani *et al.* 2019b).

The aim of this study is to assess the bivariate genomic prediction models which incorporate pairwise SNP interactions with the target of borrowing information across years to maximize the predictive ability. Since the accuracy of genomic prediction of phenotypes was shown to be increased by both borrowing information across environments and years (Covarrubias-Pazaran *et al.* 2018; Schrag *et al.* 2019b) and inclusion of epistasis into the prediction model (Martini *et al.* 2016; Vojgani *et al.* 2020), we combine these two approaches to make the best use of the available information. We further aim to assess the optimum proportion of SNP interactions to be kept in the model in the variable selection step across years. The data used for this purpose were generated in multi-location trials of doubled haploid (DH) lines generated from two European maize landraces in 2017 and 2018.

## Materials and Methods

### Data used for analysis

A set of 948 doubled haploid lines of the European maize landraces Kemater Landmais Gelb (KE, Austria, 516 lines) and Petkuser Ferdinand Rot (PE, Germany, 432 lines) were genotyped with the 600 k Affymetrix^®^ Axiom^®^ Maize Array (Unterseer *et al.* 2014).

After quality filtering and imputation, 910 DH lines remained (501 lines in KE and 409 lines in PE) and the panel of markers reduced to 501,124 markers (Hölker *et al.* 2019). Additionally, loci which were in high level of pairwise linkage disequilibrium (LD) were removed (Calus and Vandenplas 2018) through linkage disequilibrium based SNP pruning with PLINK v1.07 (Purcell *et al.* 2007; Chang *et al.* 2015). LD pruning was done by the parameters of 50, 5 and 2 which considered as the SNPs window size, the number of SNPs at which the SNP window shifts and the variance inflation factor, respectively. This resulted in a data panel containing 25’437 SNPs for KE and 30’212 SNPs for PE (Vojgani *et al.* 2020). Note that even a panel of 25’000 SNPs results in more than 1 billion SNP interactions to account for.

Out of 910 genotyped lines only 873 DH lines were phenotyped (471 lines in KE and 402 lines in PE). Einbeck (EIN, Germany), Roggenstein (ROG, Germany), Golada (GOL, Spain) and Tomeza (TOM, Spain) were the four locations that these lines were phenotyped for a series of traits in both 2017 and 2018.

The means, standard deviations, maximum and minimum values of studied phenotypic traits in 2017 and 2018 in each landrace are compared in Table 1 which were derived from the Best Linear Unbiased Estimations (BLUEs) of the genotype mean for each phenotypic trait by Hölker *et al.* (2019). The comparison of the respective detailed values for each trait in each environment and landrace in 2017 and 2018 are illustrated in the supplementary (Table S1). Vi in phenotypic traits represents the vegetative growth stage when *i* leaf collars are visible based on the leaf collar method of the corn growth (Abendroth *et al.* 2011). Early vigour at V3 stage (EV_V3), female flowering (FF) and root lodging (RL) were not phenotyped in all four environments for both years. EV_V3 was not phenotyped in EIN in 2018, FF was not phenotyped in GOL in 2017 and RL was not phenotyped in TOM and GOL in both 2017 and 2018.

**Table 1:** Phenotypic trait description and the mean, minimum, maximum and standard deviation of the BLUEs for each phenotypic trait in KE and PE landraces in the years 2017 and 2018.

| Trait | Definition | Landrace | Year | Mean | Minimum | Maximum | Standard deviation |
| --- | --- | --- | --- | --- | --- | --- | --- |
| **EV_V3** | Early vigour at V3 stage scored on scale from 1 (very poor early vigour) to 9 (very high early vigour) | KE | 2017 | 4.94 | 0.78 | 9.00 | 1.35 |
|  |  |  | 2018 | 5.06 | 0.32 | 8.67 | 1.33 |
|  |  | PE | 2017 | 5.57 | 1.00 | 9.03 | 1.20 |
|  |  |  | 2018 | 5.47 | 1.38 | 8.93 | 1.13 |
| **EV_V4** | Early vigour at V4 stage scored on scale from 1 (very poor early vigour) to 9 (very high early vigour) | KE | 2017 | 4.84 | 0.67 | 8.29 | 1.30 |
|  |  |  | 2018 | 5.08 | 0.96 | 8.65 | 1.30 |
|  |  | PE | 2017 | 5.45 | 0.93 | 8.49 | 1.15 |
|  |  |  | 2018 | 5.25 | 1.63 | 9.07 | 1.19 |
| **EV_V6** | Early vigour at V6 stage scored on scale from 1 (very poor early vigour) to 9 (very high early vigour) | KE | 2017 | 5.13 | 0.54 | 8.75 | 1.31 |
|  |  |  | 2018 | 5.54 | 1.07 | 9.60 | 1.35 |
|  |  | PE | 2017 | 5.64 | 0.84 | 8.39 | 1.12 |
|  |  |  | 2018 | 5.38 | 1.07 | 9.68 | 1.29 |
| **PH_V4** | Mean plant height of three plants of the plot at V4 stage in cm | KE | 2017 | 33.10 | 6.90 | 88.24 | 13.95 |
|  |  |  | 2018 | 42.01 | 8.48 | 89.24 | 16.47 |
|  |  | PE | 2017 | 38.01 | 11.89 | 95.30 | 14.96 |
|  |  |  | 2018 | 46.19 | 16.14 | 93.20 | 17.78 |
| **PH_V6** | Mean plant height of three plants of the plot at V6 stage in cm | KE | 2017 | 62.03 | 8.34 | 127.54 | 19.95 |
|  |  |  | 2018 | 92.27 | 21.90 | 173.66 | 21.04 |
|  |  | PE | 2017 | 69.84 | 14.78 | 130.51 | 19.26 |
|  |  |  | 2018 | 97.80 | 50.37 | 169.71 | 19.44 |
| **PH_final** | Final plant height after flowering in<br>cm | KE | 2017 | 139.10 | 49.27 | 245.00 | 27.14 |
|  |  |  | 2018 | 146.04 | 35.41 | 265.02 | 35.74 |
|  |  | PE | 2017 | 124.09 | 30.21 | 211.14 | 24.54 |
|  |  |  | 2018 | 128.08 | 35.76 | 248.43 | 35.99 |
| **FF** | Days after sowing until female<br>flowering (days until 50% of the<br>plot showed silks) | KE | 2017 | 79.72 | 62.45 | 102.02 | 6.27 |
|  |  |  | 2018 | 76.99 | 62.22 | 100.14 | 6.09 |
|  |  | PE | 2017 | 78.85 | 59.10 | 101.50 | 6.33 |
|  |  |  | 2018 | 76.70 | 60.14 | 93.96 | 6.52 |
| **RL** | Root lodging score from 1 to 9 (1 =<br>no lodging and 9= severe lodging) | KE | 2017 | 3.38 | 0.59 | 9.58 | 2.50 |
|  |  |  | 2018 | 1.42 | 0.73 | 8.52 | 0.90 |
|  |  | PE | 2017 | 2.14 | 0.03 | 9.22 | 1.74 |
|  |  |  | 2018 | 1.21 | 0.32 | 4.69 | 0.51 |

The number of phenotyped lines per year and environment for trait PH_V4, as the main trait in this study, are summarized in Table 2. For EIN and ROG a higher number of phenotyped lines were generated in 2017. On the contrary, more lines were phenotypes in GOL and TOM in 2018.

**Table 2:** Number of KE and PE lines phenotyped in each location for the years 2017 (blue numbers) and 2018 (red numbers) for trait PH_V4.

|  | EIN<br>(2017\2018) | ROG<br>(2017\2018) | GOL<br>(2017\2018) | TOM<br>(2017\2018) |
| --- | --- | --- | --- | --- |
| Phenotyped lines in KE | 462\365 | 461\365 | 211\222 | 211\222 |
| Phenotyped lines in PE | 393\365 | 390\365 | 204\240 | 204\240 |

### Statistical models for phenotype prediction

We used the bivariate statistical framework as the basis of the genomic prediction models. In this regard, GBLUP, ERRBLUP and sERRBLUP as three different methods described in Vojgani *et al.* (2020) were used for genomic prediction of phenotypes which differ in dispersion matrices representing their covariance structure of the genetic effects. GBLUP as an additive model is based on a genomic relationship matrix calculated according to VanRaden (2008). ERRBLUP (Epistatic Random Regression BLUP) as a full epistasis model is based on all pairwise SNP interactions which generates a new marker matrix considered as a marker combination matrix. The marker combination matrix is a 0, 1 matrix indicating the absence (0) or presence (1) of each marker combination for each individual. sERRBLUP (selective Epistatic Random Regression BLUP) as a selective epistasis model is based on a selected subset of SNP interactions (Vojgani *et al.* 2019b). Vojgani *et al.* (2020) proposed estimated effect variances in the training set as the selection criterion of pairwise SNP interactions due to its robustness in predictive ability specifically when only a small proportion of interactions are maintained in the model.

### Assessment of genomic prediction models

GBLUP, ERRBLUP and sERRBLUP models have been assessed via 5-fold cross validation by randomly partitioning the original sample into 5 equal size subsamples in which one subsample was considered as the test set to validate the model, and the remaining 4 subsamples were considered as a joint training set (Erbe *et al.* 2010). The 5-fold cross validation technique was utilized with 5 replicates through which the Pearson correlation between the predicted genetic values and the observed phenotypes in the test set was considered as the predictive ability in each fold of each replicate, which then was averaged across 25 replicates. In this study, predictive ability was separately assessed for KE and PE for a series of phenotypic traits in four different environments. Besides, we calculated the traits’ prediction accuracies by dividing their predictive abilities by the square-root of the respective traits’ heritabilities (Dekkers 2007) derived from all environments in both 2017 and 2018 jointly (Table S11 in the supplementary).

Univariate GBLUP within 2018 was assessed by training the model in the same year (2018) as the test set was sampled from. However, bivariate GBLUP, ERRBLUP and sERRBLUP were assessed by training the model with both the training set of the target environment in 2018 and the full dataset of the respective environment in 2017. The interaction selection step in bivariate sERRBLUP is done by first using the complete dataset of target environment in 2017 to estimate all pairwise SNP interaction effect variances. Then, an epistatic relationship matrix for all lines is constructed based on the subset of top ranked interaction effect variances, which is finally used to predict phenotypes of the target environment test set in 2018 (Vojgani *et al.* 2020).

### Variance component estimation

Variance component estimation in univariate GBLUP was done by EMMREML (Akdemir and Godfrey 2015) based on the training set in each run of 5-fold cross validation with 5 replicates. In bivariate models this was done by ASReml-R (Butler *et al.* 2018) with the approach specified by Vojgani *et al.* (2020) for pre estimating the variance components from the full dataset to derive the initial values for the variance components in ASReml models in 100 iterations for each combination. If the variance estimation based on the full set did not converge after 100 iterations, then the estimated variance components at the 100^th^ iteration were extracted as initial values of the bivariate model in the cross validation step. Afterwards, the model used these values to re-estimate the variance components based on the training set in each run of 5-fold cross validation in 50 iterations. The estimated variance components in the converged models based on the full set deviated only slightly from the estimated variance components based on the training set (Fig. 1). However, the variance component estimations did not converge in all folds of 5-fold cross validation with 5 replicates. In such cases, the initial values were set as the fixed values for the model to predict the breeding values. This approach appears justifiable in the case of non-convergence of the bivariate model, since we have shown in Fig. 2 that the difference in mean predictive ability of all folds and only the converged folds is not critical. This difference can get higher as the number of non-converged folds increases. The number of not converged folds in all studied material is shown in the supplementary (Table S12).

**Fig. 1:**
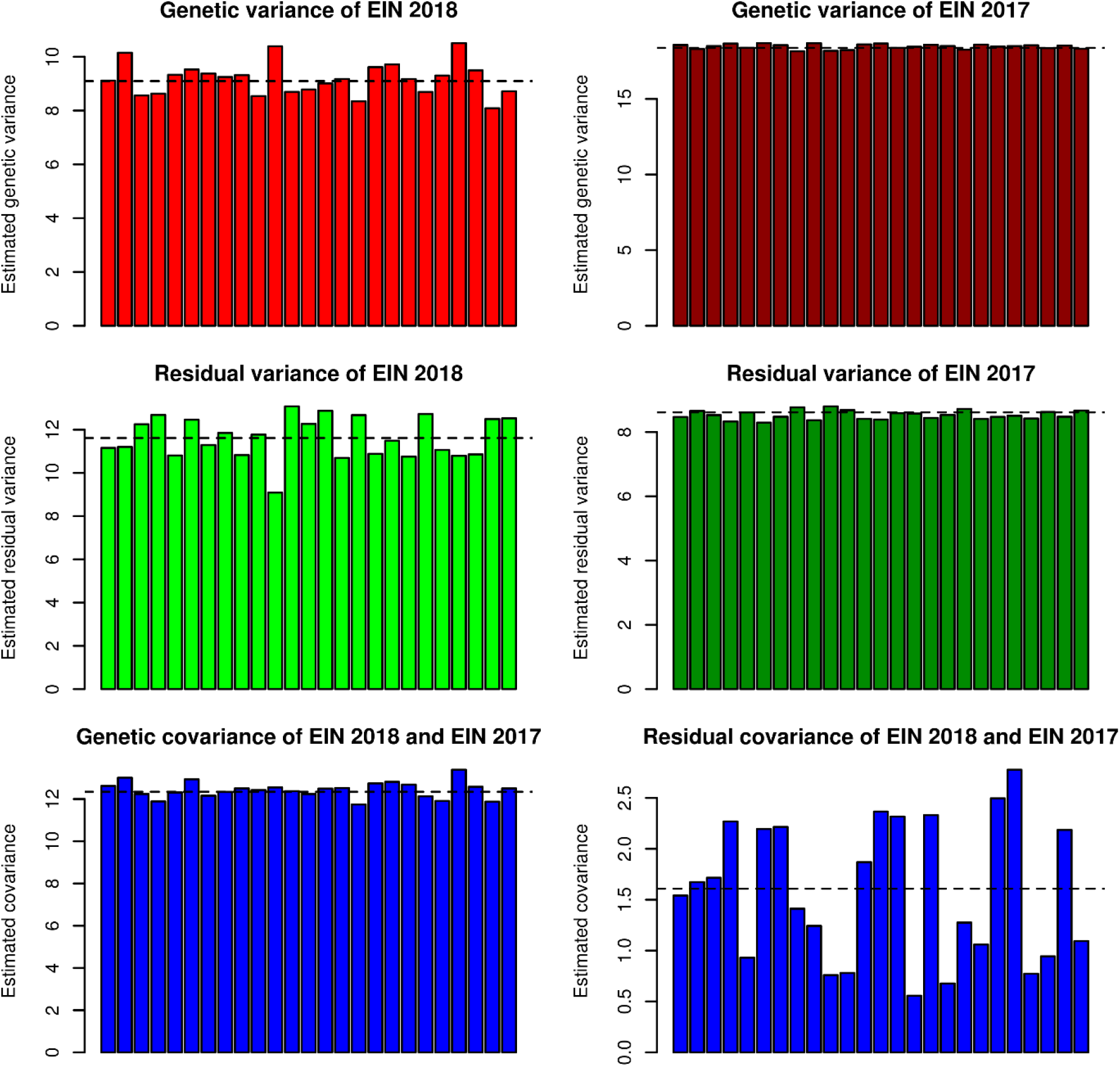
Comparison of pre estimated genetic and residual variances and covariances of converged bivariate sERRBLUP (top 10%) based on the full dataset (dashed horizontal lines) and estimated genetic and residual variances and covariances of converged bivariate sERRBLUP (top 10%) based on training set in each run of 5-fold cross validation with 5 replicates (colored bars) for predicting EIN in 2018 when the additional environment is EIN in 2017 in KE for trait PH-V4.

**Fig. 2:**
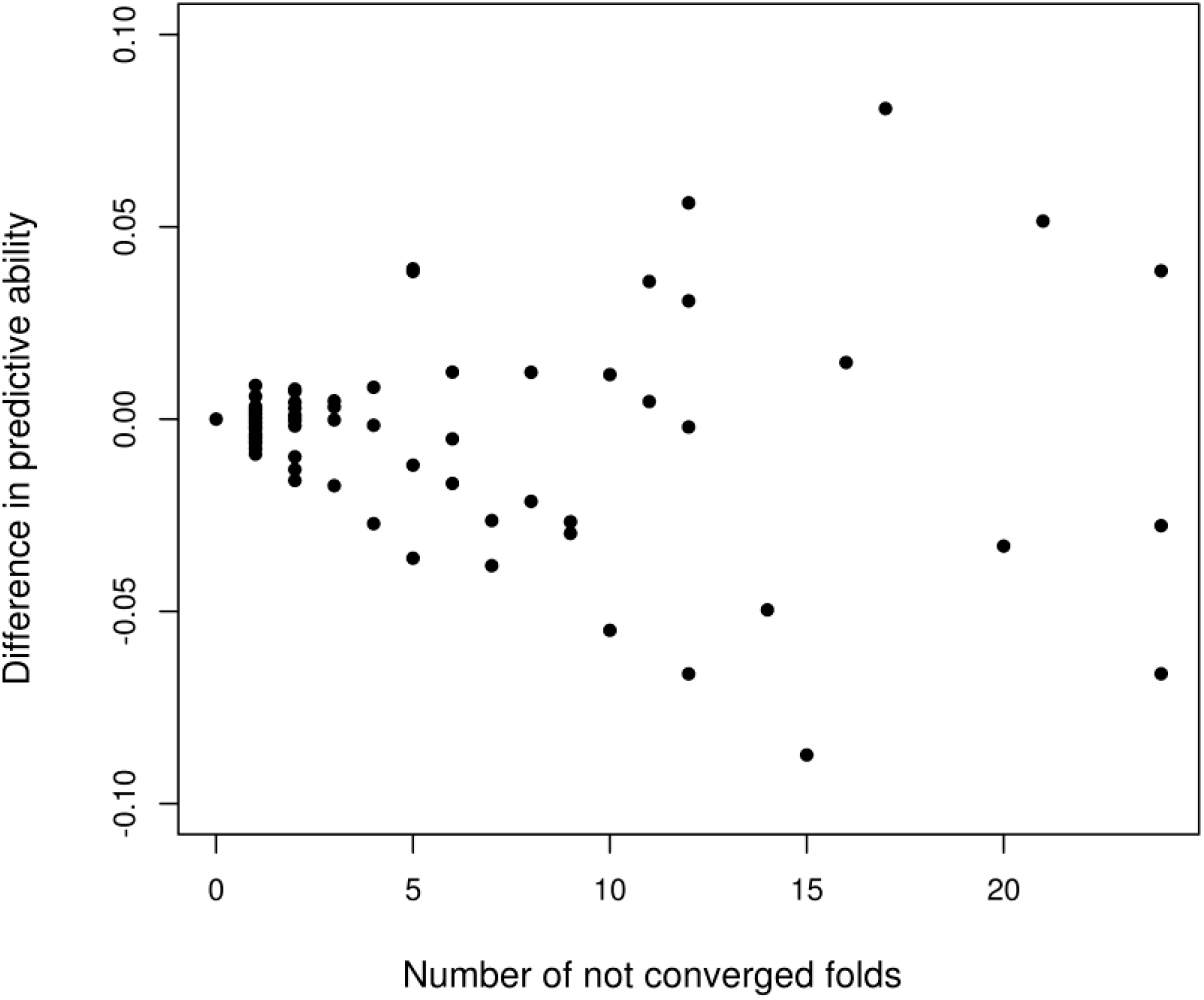
The difference between the mean predictive ability of only the converged folds and the mean predictive ability of all folds in 5-fold cross validation with 5 replicates virus the number of the folds which did not converged across all traits in all combinations for both KE and PE in bivariate GBLUP, ERRBLUP, sERRBLUP.

### Genomic correlation estimation

Genomic correlations were estimated from the genetic variances and covariance derived from the ASReml bivariate model based on the full dataset of each environment in both 2017 and 2018.

## Results

Bivariate models outperform the univariate models (Vojgani *et al.* 2020) and this has been confirmed in our study through the comparison in predictive ability of bivariate GBLUP and univariate GBLUP for the trait PH-V4 in both landraces indicating the superiority of bivariate GBLUP to univariate GBLUP in most cases (see Fig. 3). Among the bivariate genomic prediction models, bivariate ERRBLUP increases the predictive ability only slightly compared to bivariate GBLUP in a range from +0.008 to +0.024 for the trait PH-V4 across all environments in both landraces. This predictive ability increases further in bivariate sERRBLUP and the highest gain in accuracy is generally obtained when the top 10 or 5 percent of pairwise SNP interactions kept in the model in most cases. A too strict selection like using only the top 0.001 percent interactions, results in a decrease in predictive ability (see Fig. 3). Robustness of the predictive ability depending on the share of selected markers was higher in PE. Similar patterns are observed across a series of other traits for bivariate models which are shown in the supplementary (Fig. S1-S7). Additionally, the predictive ability of univariate GBLUP by training the model on the average phenotypic values of both 2017 and 2018 was evaluated for a series of phenotypic traits, which yielded quite similar predictive ability as obtained with univariate GBLUP within year 2018 or worse in some cases (Table S10a (KE) and S10b (PE) in supplementary).

**Fig. 3:**
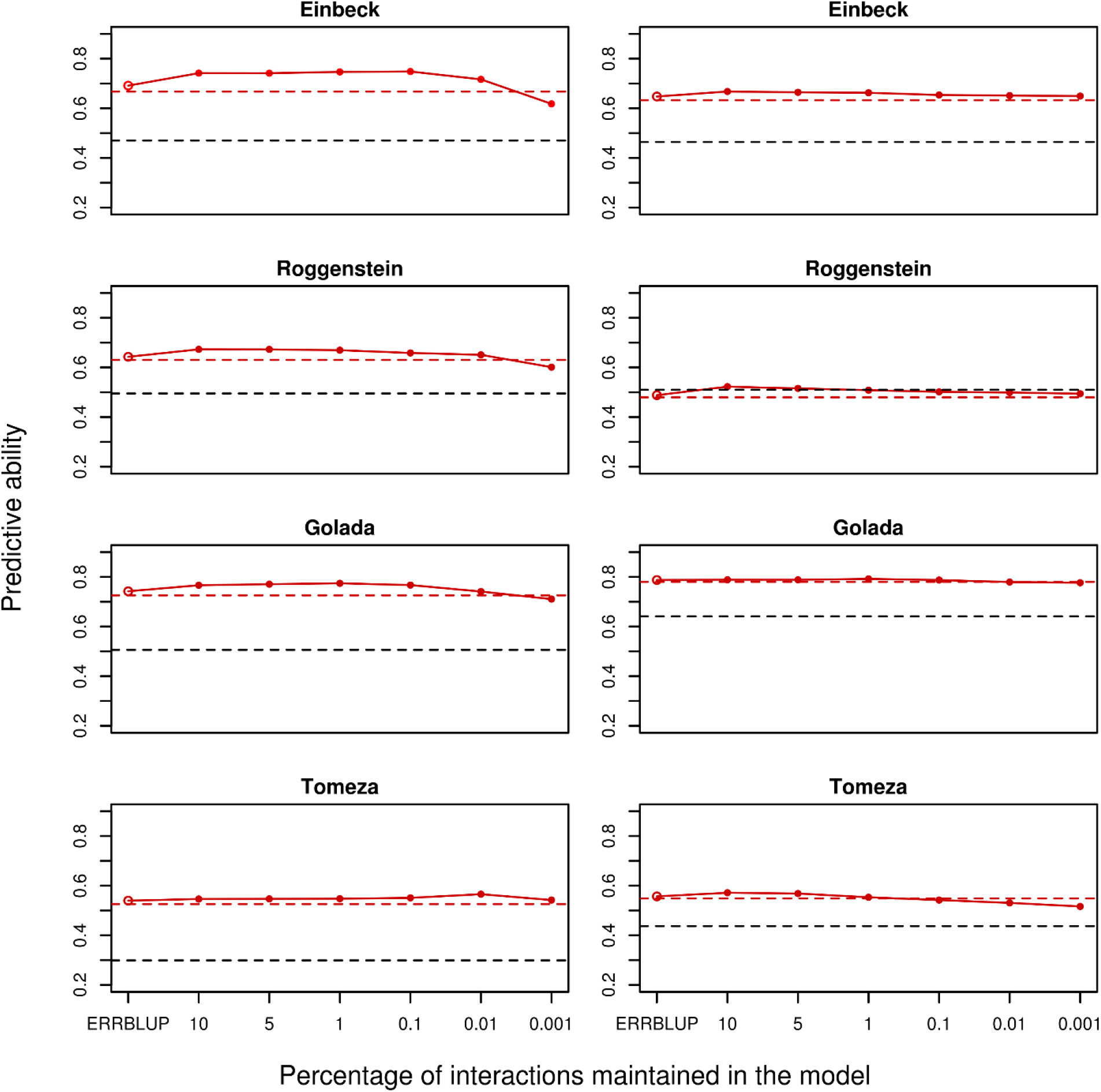
Predictive ability for univariate GBLUP within 2018 (black dashed horizontal line), bivariate GBLUP (red dashed horizontal line), bivariate ERRBLUP (red open circle) and bivariate sERRBLUP (red filled circles and red solid line) for trait PH-V4 in KE (left) and in PE (right).

The absolute gain in predictive ability from univariate GBLUP to maximum bivariate sERRBLUP was regressed on the respective sERRBLUP genomic correlation between the two respective environment and across the series of studied traits (Fig. 4). Regression coefficients range between 0.09 and 0.51 and thus show a clear association between the absolute gain in prediction accuracy and the genomic correlation between environments. When combining all traits and environments, this correlation is 0.64 (p-value = 0.00024) in KE and 0.73 (p-value = 1.072e-05) in PE.

**Fig. 4:**
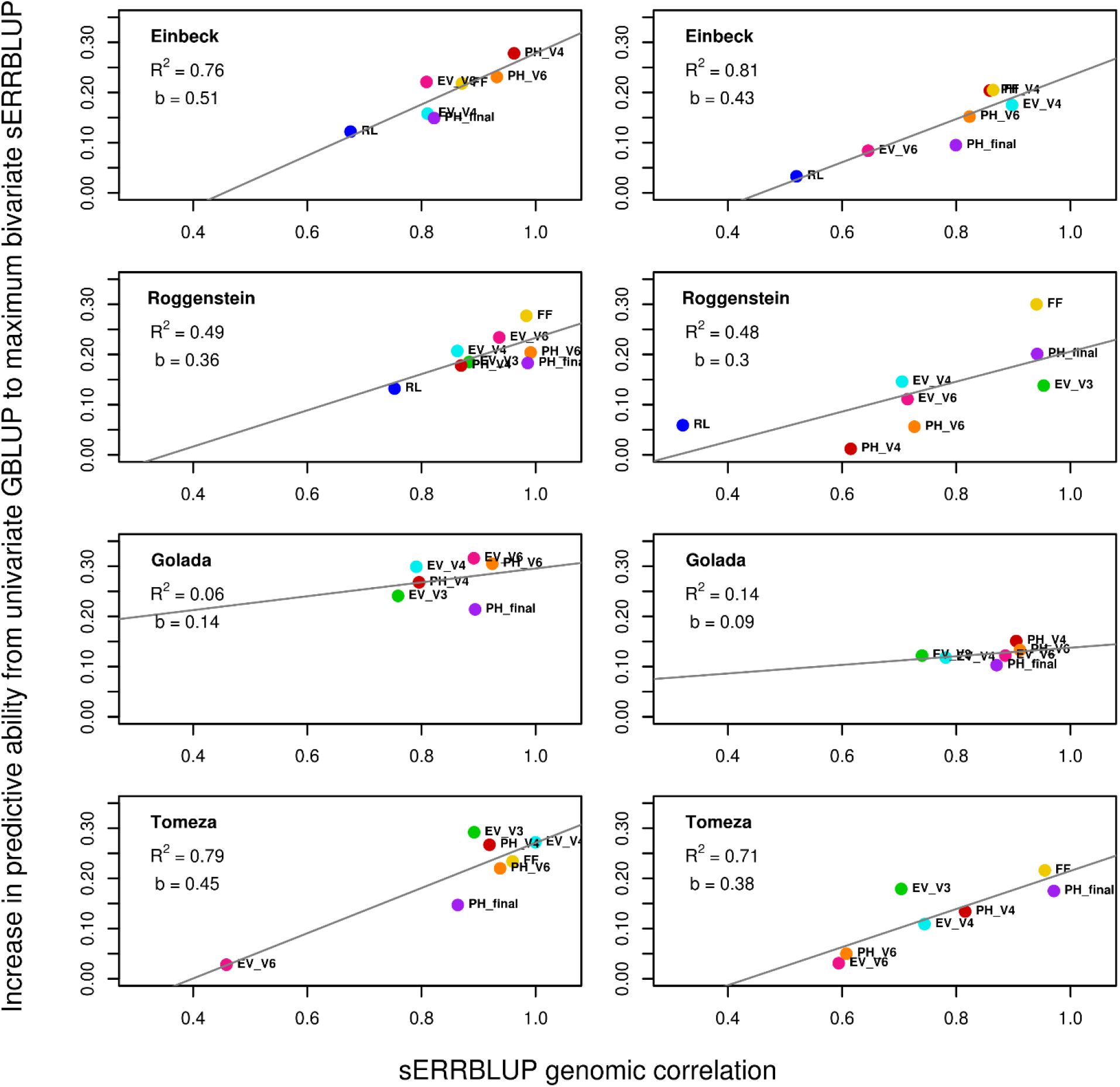
Regression of the absolute increase in predictive ability from univariate GBLUP to maximum bivariate sERRBLUP on the respective sERRBLUP genomic correlation between 2017 and 2018 in KE (left) and in PE (right) for all studied traits. In each panel, the overall linear regression line (gray solid line) with the regression coefficient (**b**) and R-squared (**R**^2^) are shown.

The genomic correlations across years estimated with GBLUP and sERRBLUP for the trait PH_V4 are illustrated in Table 3, indicating that the proportion of interactions in bivariate sERRBLUP which maximized the predictive ability are not necessarily linked to the highest genomic correlation. In contrast, the best sERRBLUP for trait PH_V4 is linked to the lowest genomic correlation in most cases. However, this is not the general pattern observed for series of other traits and the best sERRBLUP for some traits and environments combinations are linked to the highest genomic correlation (Table S3-S9 in supplementary).

**Table 3:**
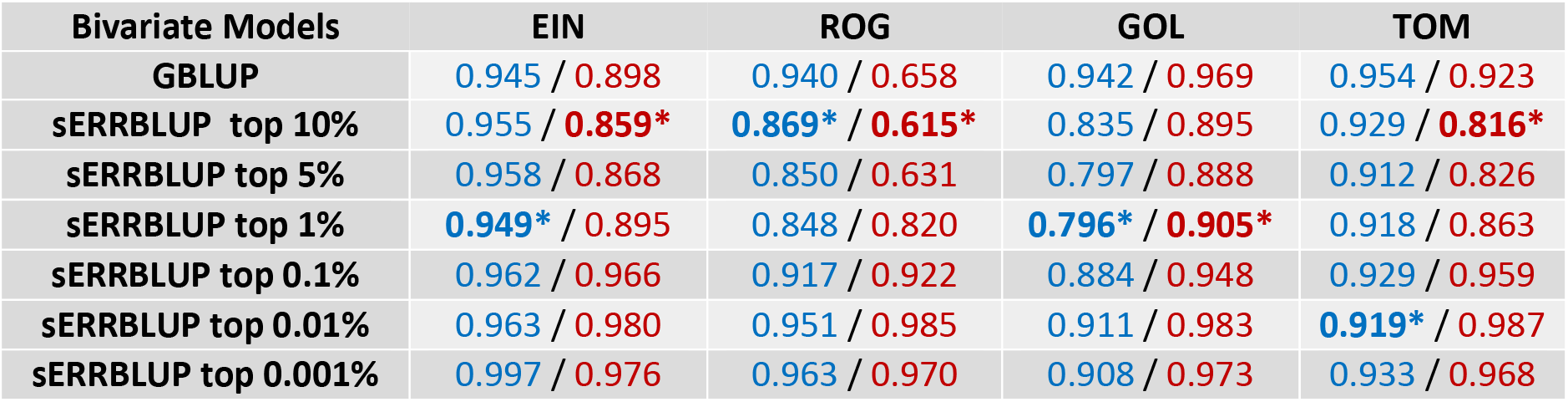
Genomic correlation between 2017 and 2018 in each environment for trait PH_V4 for KE (blue numbers) and PE (red numbers). The blue and red bold numbers with stars indicate which proportion of interactions in bivariate sERRBLUP maximized the predictive ability in each environment for KE and PE, respectively.

| Bivariate Models | EIN | ROG | GOL | TOM |
| --- | --- | --- | --- | --- |
| GBLUP | 0.945 / 0.898 | 0.940 / 0.658 | 0.942 / 0.969 | 0.954 / 0.923 |
| sERRBLUP top 10% | 0.955 / **0.859\*** | **0.869\*** / **0.615\*** | 0.835 / 0.895 | 0.929 / **0.816\*** |
| sERRBLUP top 5% | 0.958 / 0.868 | 0.850 / 0.631 | 0.797 / 0.888 | 0.912 / 0.826 |
| sERRBLUP top 1% | **0.949\*** / 0.895 | 0.848 / 0.820 | **0.796\*** / **0.905\*** | 0.918 / 0.863 |
| sERRBLUP top 0.1% | 0.962 / 0.966 | 0.917 / 0.922 | 0.884 / 0.948 | 0.929 / 0.959 |
| sERRBLUP top 0.01% | 0.963 / 0.980 | 0.951 / 0.985 | 0.911 / 0.983 | **0.919\*** / 0.987 |
| sERRBLUP top 0.001% | 0.997 / 0.976 | 0.963 / 0.970 | 0.908 / 0.973 | 0.933 / 0.968 |

In this regard, the absolute increase in predictive ability from bivariate GBLUP to maximum bivariate sERRBLUP was regressed on the difference between genetic correlations estimated with GBLUP and maximum sERRBLUP, respectively, across all traits in both landraces. Fig. 5 shows a significant correlation of 0.42 (p-value = 0.0255) in KE and 0.74 (p-value = 6.458e-06) in PE between the absolute gain in the respective predictive ability and the difference in the corresponding genetic correlations.

**Fig. 5:**
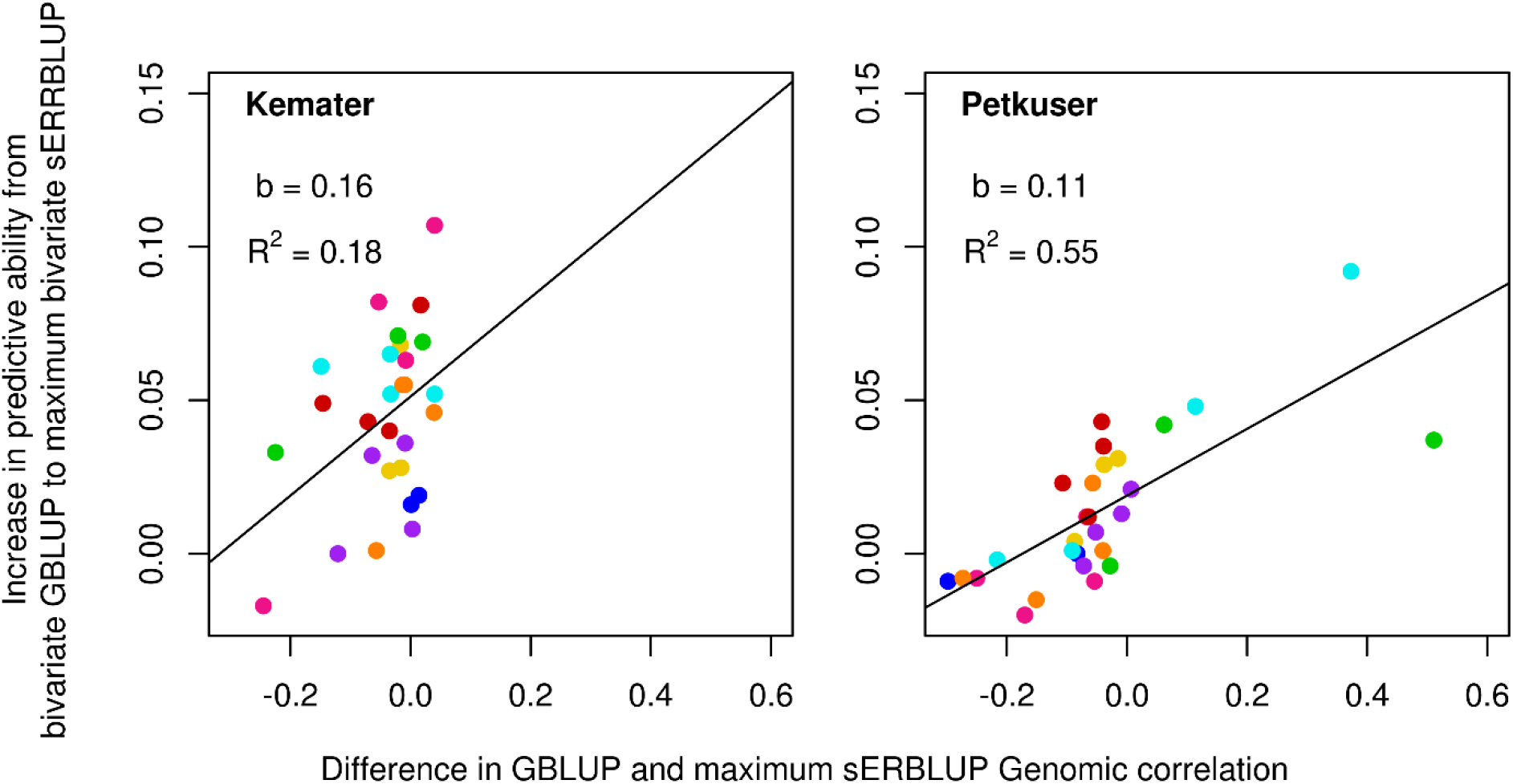
Regression of the absolute increase in predictive ability from bivariate GBLUP to maximum bivariate sERRBLUP on the difference between the GBLUP genomic correlation and maximum sERRBLUP genomic correlation between 2017 and 2018 in KE (left) and in PE (right) for all studied traits. In each panel, the overall linear regression line with the regression coefficient (**b**) and R-squared (**R**^2^) are shown. The colors green, light blue, pink, red, orange, purple, yellow and dark blue represent the phenotypic traits EV_V3, EV_V4, EV_V6, PH_V4, PH_V6, PH_final, FF and RL, respectively.

There might be some tendency that including phenotypes of the previous year into prediction becomes more efficient when the phenotypic correlation between years is high. In this context, the correlation between the absolute gain in predictive ability from univariate GBLUP to maximum bivariate sERRBLUP and the phenotypic correlation among the years (see Table S2) over all studied traits in all four environments and in both landraces was studied. Fig. 6 demonstrates that the maximum correlation between the absolute gain in the respective predictive ability and the phenotypic correlation is obtained in EIN for KE (0.69) and in TOM for PE (0.72). Across all studied traits and environments, there is a significant correlation of 0.59 in KE (p-value= 0.001) and 0.47 in PE (p-value= 0.01).

**Fig. 6:**
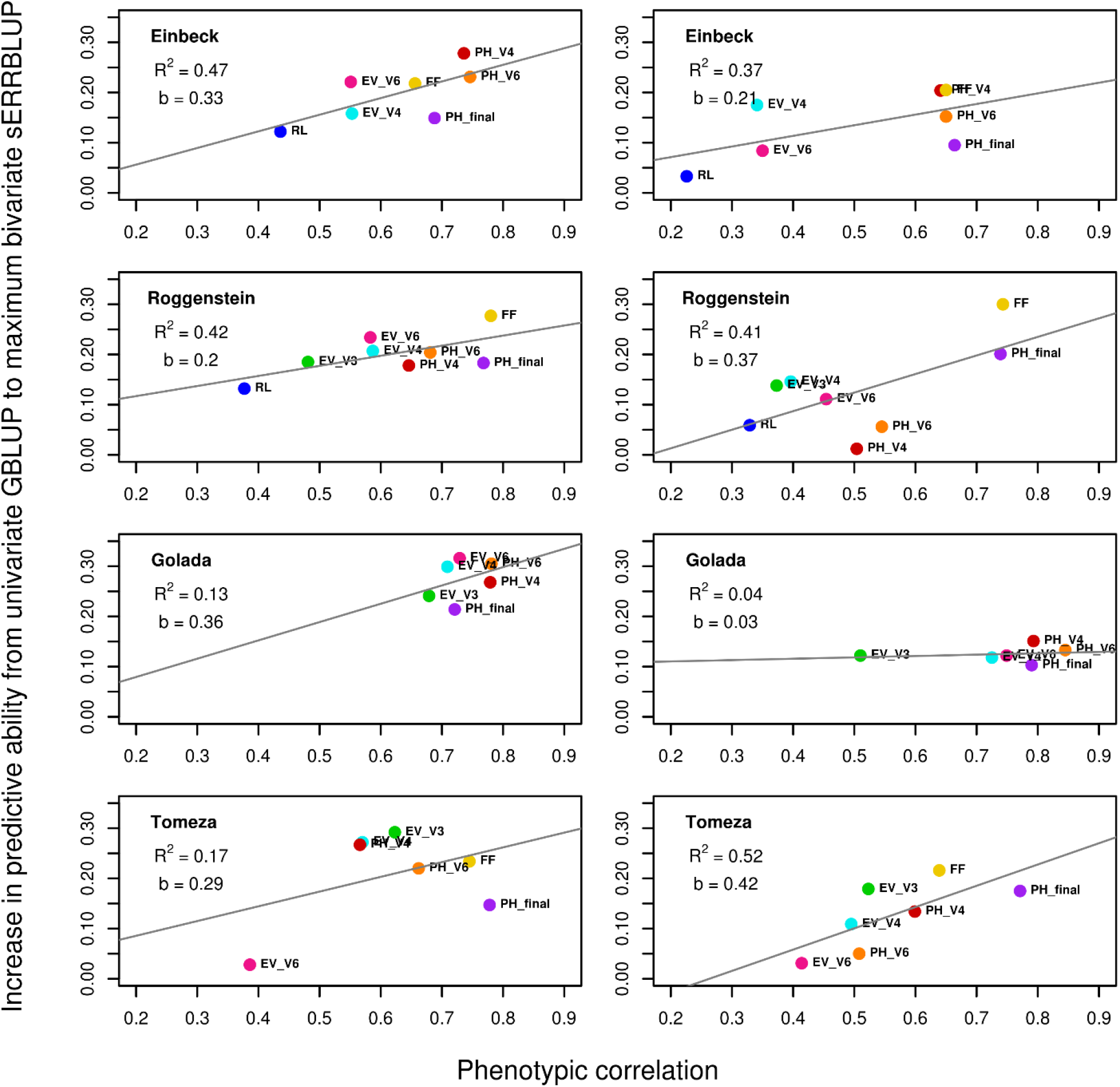
Regression of the absolute increase in predictive ability from univariate GBLUP to maximum bivariate sERRBLUP on the phenotypic correlation between 2017 and 2018 in KE (left) and in PE (right) for all studied traits. In each panel, the overall linear regression line (gray solid line) with the regression coefficient (**b**) and R-squared (**R**^2^) are shown.

Overall, the percentage of relative increase in prediction accuracy from the bivariate GBLUP to the maximum bivariate sERRBLUP in both landraces reveals more increase in prediction accuracy for KE than PE with the average increase of 7.61 percent in KE and 3.47 percent in PE over all studied traits (see Fig. 7). Among all traits, the maximum increase in prediction accuracy for KE is 22.63 percent which was obtained in EV_V6 in EIN, and for PE is 34.59 percent which was obtained in EV_V4 in EIN. However, Fig. 7 shows some slight decreases in prediction accuracy from bivariate GBLUP to maximum bivariate sERRBLUP for some combinations of traits and environment in both landraces. This is more often observed in PE than KE, where the maximum decrease was found in EV_V6 in TOM for both PE (−3.198 percent) and KE (−2.795 percent). Overall, the average relative increase from bivariate GBLUP to maximum bivariate sERRBLUP was over 3 percent in most cases. The absolute increase in prediction accuracy is also illustrated in the supplementary (Fig. S8) indicating the average increase of 0.046 in KE and 0.015 in PE over all combinations of traits and environments.

**Fig. 7:**
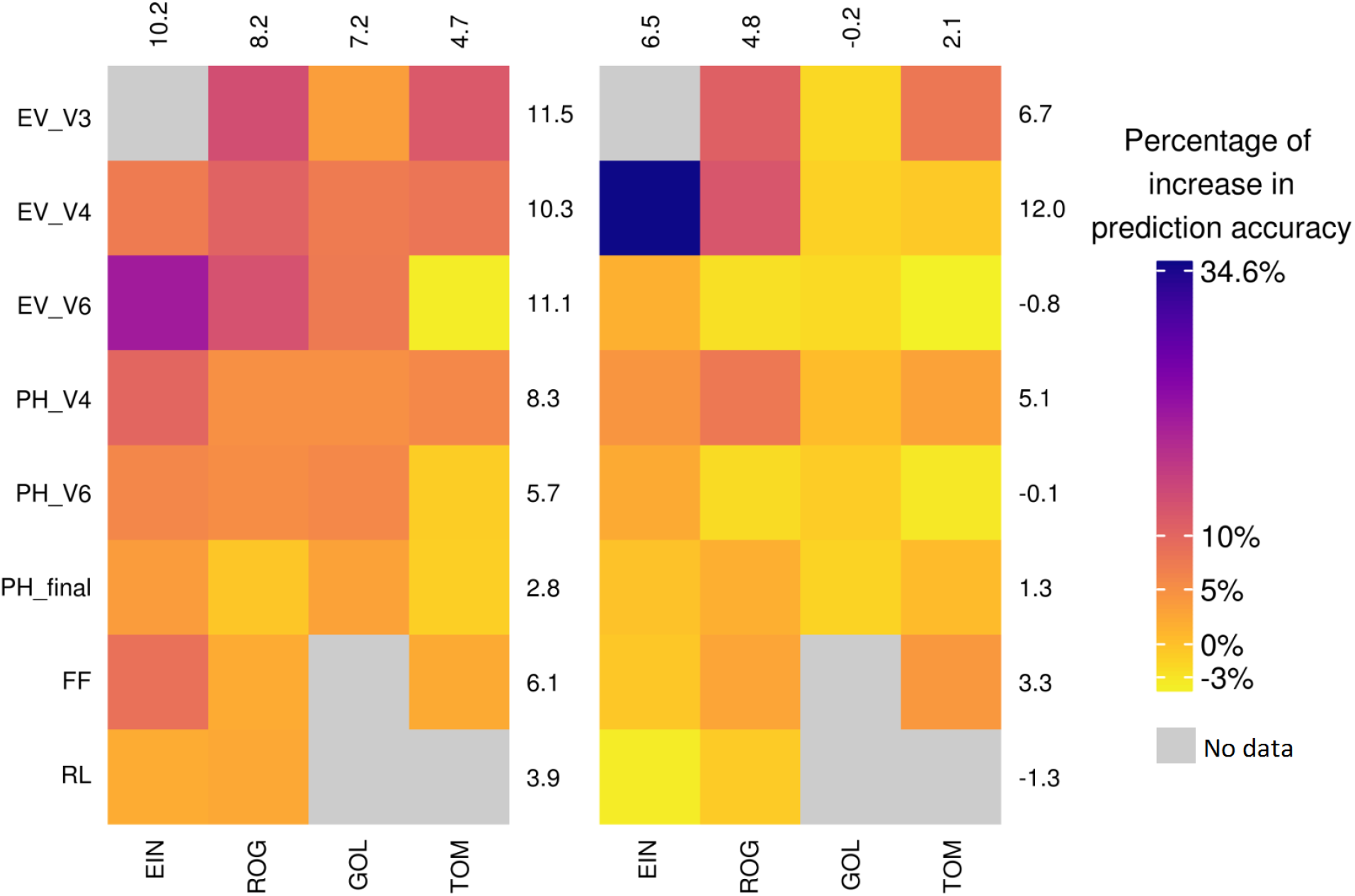
Percentage of change in prediction accuracy from bivariate GBLUP to the maximum prediction accuracy of bivariate sERRBLUP in KE (left side plot) and in PE (right side plot). The average percentage of change in prediction accuracy for each trait and environment is displayed in all rows and columns, respectively.

Finally, a comparison between the absolute increase in prediction accuracy from bivariate GBLUP to maximum bivariate sERRBLUP in PE versus KE shows a higher increase in KE compared to PE with a regression coefficient 0.25 (see Fig. 8). This indicates some consistency of the observed trends across landraces. This was also confirmed with paired t-test indicating that the mean increase in prediction accuracy for KE is significantly higher than in PE (p-value= 3.921e-05).

**Fig. 8:**
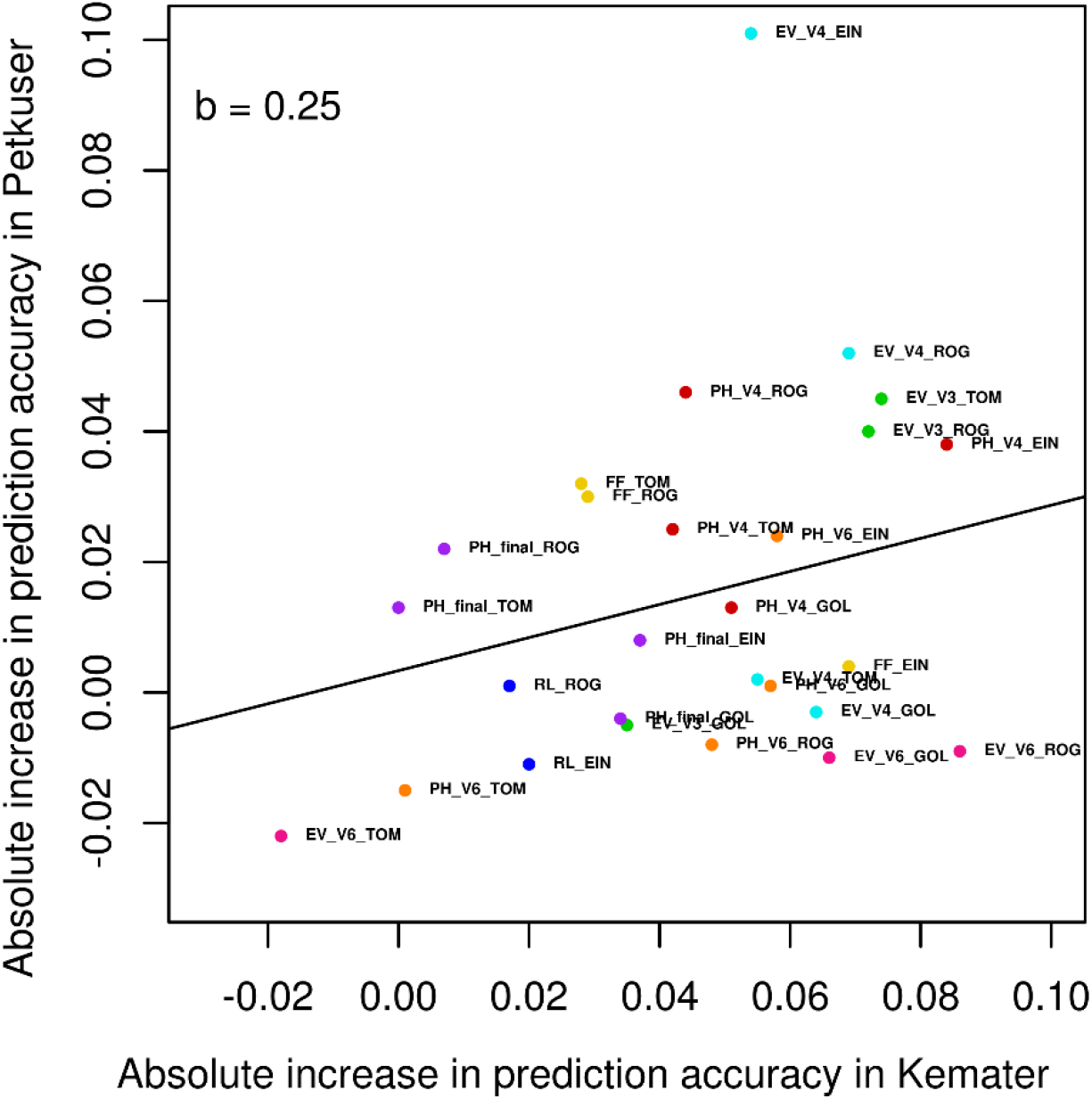
Absolute change in prediction accuracy from bivariate GBLUP to the maximum prediction accuracy of bivariate sERRBLUP in PE vs. KE. The black line represents the overall linear regression line.

## Discussion

In this study, bivariate ERRBLUP as a full epistasis model incorporating all pairwise SNP interactions provides only a modest increase in predictive ability compared to bivariate GBLUP. This was expected, since ERRBLUP incorporates a high number of interactions by which a large number of unimportant variables are introduced into the model (Martini *et al.* 2016), thus introducing potential ‘noise’ which can prevent gains in predictive ability. In contrast, bivariate sERRBLUP substantially increases the predictive ability compared to bivariate GBLUP. In fact, the increase in predictive ability from bivariate GBLUP to bivariate sERRBLUP is only caused by inclusion of relevant pairwise SNP interactions. Note that all bivariate models substantially outperformed univariate GBLUP, as phenotypic data of the respective environment in the previous year was used.

It was shown that multivariate GBLUP is superior in predictive ability compared to univariate GBLUP under existence of medium (~0.6) or high (~0.9) genomic correlation, and that the low genomic correlation results in no increase in multivariate GBLUP compared to univariate GBLUP (Covarrubias-Pazaran *et al.* 2018). Calus *et al.* (2011) also found an increase of 3 to 14 percent in predictive ability of multi-trait SNP-based models in a simulation study when genetic correlations ranged from 0.25 to 0.75. In our study, we also found a significant correlation between the absolute gain in prediction accuracy from univariate GBLUP to maximum bivariate sERRBLUP and the respective genomic correlation in both KE (r = 0.64) and PE *(r*= 0.73) across all traits and environments combinations.

Moreover, Martini *et al.* (2016) showed that the predictive ability in one environment can be increased by variable selection in the other environment under the assumption of positive phenotypic correlation between environments. It was shown in a wheat dataset (Pérez and de los Campos 2014), where environments 2 and 3 had the highest phenotypic correlation (0.661), that the predictive ability for phenotype prediction in environment 2 was maximized by variable selection in environment 3 and vice versa (Martini *et al.* 2016). Therefore, the increase in prediction accuracy is expected to be influenced by the phenotypic correlations between the environments or between the years in the same environment in bivariate models. In our study, although 2017 and 2018 were climatically quite different, since 2018 suffered from a major heat stress compared to 2017 (Table 1), we see a significant correlation between the absolute gain in predictive ability from univariate GBLUP to maximum predictive ability of bivariate sERRBLUP and the phenotypic correlation between years in each environment for both KE *(r =* 0.59) and PE (r = 0.47).

In addition to the genomic and phenotypic correlations between the years, the trait heritability is another factor which is expected to be influential for such an increase in bivariate sERRBLUP predictive ability as well. Therefore, the traits with lower heritability are expected to obtain less gain in sERRBLUP predictive ability than the traits with higher heritability. In our study, the correlation between the absolute gain in prediction accuracy from univariate GBLUP to maximum bivariate sERRBLUP and a trait’s heritability over all studied material was considerable in both KE (r = 0.35) and PE (r = 0.45) (Fig. S9 in the supplementary). Based on the obtained results, the traits with low heritability (e.g. 0.59 for RL in PE) showed only a small increase in prediction accuracy. However, not all traits with higher heritabilities did necessarily show a higher gain in predictive ability for all traits. Overall, this association between the absolute gain in predictive ability and the trait heritabilities were close to significant in KE (p-value=0.07) and highly significant in PE (p-value=0.02). It should be noted that the trait heritabilities were calculated on an entry-mean basis within each KE and PE landraces (Hallauer *et al.* 2010) over all eight environments in both years 2017 and 2018 jointly. The trait heritabilities obtained only from 2017 are significantly higher than the trait heritabilities obtained only from 2018 in both KE and PE based on a paired t-test (Table S11 in the supplementary). This also results in an increase in predictive ability from univariate GBLUP to maximum bivariate sERRBLUP in KE and PE, since multi-trait models have the potential of increasing the predictive ability when traits with low heritability are joined with traits with higher heritability, given they are genomically correlated (Thompson and Meyer 1986).

It should be noted that the increase in predictive ability from univariate GBLUP to maximum bivariate sERRBLUP is caused by both borrowing information across years and capitalizing on epistasis, while the increase in predictive ability from bivariate GBLUP to maximum bivariate sERRBLUP is caused by accounting for epistasis alone. Overall, the traits behave differently among different environments and landraces due to their genomic correlations, phenotypic correlations and heritabilities. To shed light on this, the maximum increase in prediction accuracy from bivariate GBLUP to bivariate sERRBLUP in KE was observed for the trait EV_V6 (0.112) in EIN where the corresponding sERRBLUP genomic correlation (0.809) is higher than the GBLUP genomic correlation (0.768). This trait has a high heritability (0.90) and high phenotypic correlation (0.551) as well. In contrast, the respective prediction accuracy decreases (−0.018) for EV_V6 in TOM for KE indicating the lower sERRBLUP genomic correlation (0.458) than GBLUP genomic correlation (0.703) and the particularly low phenotypic correlation (0.383). It should be noted that the phenotypic correlation does not play a major role for the increase in prediction accuracy from bivariate GBLUP to bivariate sERRBLUP, since both models are bivariate and benefit from the same phenotypic correlations. Therefore, EV_V6 obtaining the maximum and minimum increase in the respective prediction accuracy for KE indicates the significant role of genomic correlation among the possible causes. In general, bivariate sERRBLUP improves the prediction accuracy compared to bivariate GBLUP more in KE than PE which is potentially due to significantly higher sERRBLUP genomic correlation and heritability in KE compared to PE, based on paired t-test.

In our study, 5-fold cross validation with 5 replicates was utilized to evaluate our bivariate genomic prediction models. Different split of cross validation such as 10-fold cross validation did not make a considerable difference in our bivariate models’ predictive abilities (Fig. S10 in the supplementary). The maximum increase in bivariate models’ predictive abilities when utilizing 10-fold cross validation with 10 replicates compared to utilizing 5-fold cross validation with 5 replicates was 0.018 in KE and 0.006 in PE for trait PH_V4. Overall, our cross validation scenario is not expected to bias the predictive abilities obtained from our bivariate models for reasons as outlined by Runcie and Cheng (2019), who observed a bias when the test set of the target trait is predicted from the full dataset of the second trait in multi-trait model. In our study, utilizing the full dataset of the target trait in one environment from 2017 to predict the same biological trait in the respective environment in 2018 should not lead to such a bias in predictive ability, since the individuals do not share the same source of non-genetic variation and they have been grown in two different years which have been climatically very different from each other.

Overall, our results indicate that incorporating a suitable subset of epistatic interactions besides utilizing information across years can substantially increase the predictive ability. The amount of this increase is affected by the genomic and phenotypic correlations between the years and the heritability of the phenotypic trait. Therefore, this approach is potentially beneficial for genomic prediction of phenotypes under the assumption of sufficient genomic and phenotypic correlation between years for highly heritable traits. This may allow to reduce the number of lines which have to be phenotyped over several years and thus reduce phenotyping costs which and thus be of high interest in practical plant breeding.

## Declaration

### Ethics approval

The authors declare that this study complies with the current laws of the countries in which the experiments were performed.

### Consent to participate

Not applicable

### Consent for publication

Not applicable

### Availability of data and materials

All data and material are available through material transfer agreements upon request.

### Code availability

Not applicable

### Authors’ contributions

EV derived the results, analyzed the data, wrote the manuscript; TP proposed epistasis relationship matrices; ACH, MM and CCS prepared the material; ACH proposed cross validation strategy in bivariate model; HS proposed the original research question, guided the structure of the research.TP ACM MM CCS HS read, revised and approved the manuscript.

## Acknowledgements

We are thankful to KWS SAAT SE, Misión Biológica de Galicia, Spanish National Research Council (CSIC), Technical University of Munich, and University of Hohenheim for providing the extensive phenotypic evaluation. We are grateful to the German Federal Ministry of Education and Research (BMBF) for the funding of our project within the scope of the funding initiative “Plant Breeding Research for the Bioeconomy” (MAZE – “Accessing the genomic and functional diversity of maize to improve quantitative traits”; Funding ID: 031B0195).

