## Supplementary Material for "Bivariate genomic prediction of phenotypes by selecting epistatic interactions across years"

Theoretical and Applied Genetics

**Supplemental Tables**

**Table S1** The mean, minimum, maximum and standard deviation of BLUEs of phenotypic traits in each location for KE (blue numbers) and PE (red numbers) in 2017 and 2018.

| Trait | Location/Year | Mean | Minimum | Maximum | Standard deviation |
|---|---|---|---|---|---|
| EV_V3 | ROG/2017 | 5.35\5.84 | 1.71\2.90 | 7.90\7.92 | 0.95\0.75 |
| | ROG/2018 | 5.68\5.91 | 1.21\3.46 | 7.91\8.08 | 1.01\0.89 |
| | GOL/2017 | 6.31\6.67 | 4.07\5.49 | 8.49\7.98 | 0.69\0.51 |
| | GOL/2018 | 4.26\4.62 | 0.32\1.69 | 7.63\7.99 | 1.20\0.97 |
| | TOM/2017 | 5.51\6.15 | 1.93\3.84 | 7.34\8.45 | 0.99\0.67 |
| | TOM/2018 | 4.84\5.67 | 0.59\1.38 | 8.67\8.93 | 1.43\1.13 |
| EV_V4 | EIN/2017 | 4.24\4.82 | 0.94\1.52 | 7.07\7.46 | 1.11\0.98 |
| | EIN/2018 | 4.56\4.61 | 0.98\1.99 | 7.01\6.99 | 1.05\0.78 |
| | ROG/2017 | 5.44\5.85 | 2.65\2.88 | 7.86\7.94 | 0.92\0.78 |
| | ROG/2018 | 5.61\5.84 | 1.39\2.89 | 8.41\9.07 | 1.22\1.15 |
| | GOL/2017 | 5.71\5.98 | 3.37\3.91 | 7.89\7.89 | 0.81\0.83 |
| | GOL/2018 | 5.16\5.29 | 1.50\1.63 | 8.36\8.44 | 1.26\1.36 |
| | TOM/2017 | 5.26\5.75 | 2.59\3.92 | 6.89\7.35 | 0.83\0.61 |
| | TOM/2018 | 5.01\5.27 | 0.96\2.08 | 8.56\8.07 | 1.49\1.11 |
| EV_V6 | EIN/2017 | 5.03\5.54 | 0.97\1.51 | 8.05\8.39 | 1.24\1.06 |
| | EIN/2018 | 4.73\4.73 | 1.07\2.58 | 6.95\6.08 | 0.78\0.56 |
| | ROG/2017 | 5.55\5.91 | 1.02\2.52 | 8.07\7.76 | 0.95\0.77 |
| | ROG/2018 | 6.14\6.42 | 2.21\3.36 | 8.81\9.68 | 1.29\1.20 |
| | GOL/2017 | 6.24\6.24 | 3.90\3.81 | 8.45\7.94 | 0.85\0.85 |
| | GOL/2018 | 5.12\4.77 | 1.21\1.17 | 8.23\7.51 | 1.29\1.26 |
| | TOM/2017 | 5.58\5.86 | 2.96\3.90 | 7.66\7.91 | 0.92\0.68 |
| | TOM/2018 | 6.30\5.43 | 2.44\1.07 | 9.60\9.08 | 1.36\1.26 |
| PH_V4 | EIN/2017 | 34.49\38.73 | 6.90\20.43 | 53.14\57.94 | 7.24\6.17 |
| | EIN/2018 | 32.54\35.23 | 8.48\19.60 | 49.63\50.29 | 5.65\5.24 |
| | ROG/2017 | 25.50\28.10 | 9.23\13.63 | 42.29\41.54 | 4.60\4.53 |
| | ROG/2018 | 29.08\31.75 | 11.11\17.45 | 43.70\45.04 | 4.79\4.92 |

| | | | | | |
|---|---|---|---|---|---|
| | **GOL/2017** | 62.88\68.98 | 34.30\38.39 | 88.24\95.30 | 9.79\10.96 |
| | **GOL/2018** | 60.37\64.49 | 23.27\16.14 | 89.24\93.20 | 12.54\14.32 |
| | **TOM/2017** | 41.60\47.45 | 11.98\25.37 | 63.89\72.12 | 8.71\8.27 |
| | **TOM/2018** | 60.32\66.49 | 28.43\47.45 | 84.24\88.27 | 9.59\8.03 |
| **PH_V6** | **EIN/2017** | 62.40\69.36 | 21.41\36.53 | 95.54\98.80 | 11.89\9.62 |
| | **EIN/2018** | 78.81\85.14 | 21.90\52.02 | 105.05\115.61 | 10.53\10.07 |
| | **ROG/2017** | 61.46\68.91 | 32.17\30.35 | 89.74\94.77 | 9.34\9.52 |
| | **ROG/2018** | 82.64\90.89 | 41.48\57.90 | 118.69\123.27 | 11.00\11.17 |
| | **GOL/2017** | 94.21\98.30 | 37.28\54.75 | 127.54\130.51 | 15.05\15.29 |
| | **GOL/2018** | 101.90\104.82 | 53.69\50.37 | 137.67\146.02 | 15.42\18.24 |
| | **TOM/2017** | 83.86\92.35 | 48.46\57.81 | 119.07\124.98 | 14.41\12.79 |
| | **TOM/2018** | 120.46\120.57 | 68.48\58.96 | 173.66\169.71 | 19.56\18.63 |
| **PH_final** | **EIN/2017** | 159.18\141.35 | 100.84\69.01 | 228.96\211.14 | 21.57\21.10 |
| | **EIN/2018** | 114.90\93.35 | 82.28\49.97 | 172.12\136.25 | 16.46\16.27 |
| | **ROG/2017** | 137.04\122.25 | 74.25\63.56 | 211.14\201.92 | 22.32\20.56 |
| | **ROG/2018** | 163.71\142.70 | 103.82\70.16 | 249.35\208.81 | 25.52\23.66 |
| | **GOL/2017** | 115.68\102.69 | 49.27\30.21 | 167.58\149.14 | 21.73\23.59 |
| | **GOL/2018** | 129.94\117.16 | 35.41\35.76 | 186.09\173.10 | 26.35\27.58 |
| | **TOM/2017** | 157.99\144.61 | 81.92\79.28 | 245.00\195.36 | 24.82\18.95 |
| | **TOM/2018** | 184.54\169.57 | 115.10\118.15 | 265.02\248.43 | 26.34\22.77 |
| **FF** | **EIN/2017** | 82.55\81.78 | 70.36\68.86 | 102.02\101.50 | 5.23\5.17 |
| | **EIN/2018** | 78.80\79.59 | 63.37\68.12 | 94.35\93.96 | 5.44\5.40 |
| | **ROG/2017** | 73.06\71.91 | 62.45\59.10 | 91.22\88.03 | 4.82\4.47 |
| | **ROG/2018** | 79.16\79.05 | 66.74\67.74 | 100.14\92.87 | 4.72\4.41 |
| | **TOM/2017** | 76.88\74.16 | 63.93\62.13 | 93.28\92.17 | 5.58\4.64 |
| | **TOM/2018** | 70.31\68.76 | 62.22\60.14 | 83.64\90.06 | 4.11\3.73 |
| **RL** | **EIN/2017** | 3.48\2.23 | 0.63\0.76 | 9.21\8.08 | 2.29\1.54 |
| | **EIN/2018** | 1.58\1.25 | 0.73\0.32 | 8.52\4.69 | 1.10\0.59 |
| | **ROG/2017** | 2.39\1.50 | 0.96\0.95 | 9.01\8.50 | 2.21\1.13 |
| | **ROG/2018** | 1.27\1.17 | 0.95\0.95 | 7.01\3.52 | 0.61\0.41 |

**Table S2** Phenotypic correlation between 2017 and 2018 in each environment for KE and PE.

| Trait | Landrace | EIN | ROG | GOL | TOM |
|---|---|---|---|---|---|
| **EV_V3** | **KE**<br>**PE** | -<br>- | 0.481<br>0.373 | 0.679<br>0.510 | 0.623<br>0.523 |
| **EV_V4** | **KE**<br>**PE** | 0.553<br>0.341 | 0.587<br>0.396 | 0.709<br>0.725 | 0.570<br>0.495 |
| **EV_V6** | **KE**<br>**PE** | 0.551<br>0.350 | 0.583<br>0.454 | 0.729<br>0.749 | 0.383<br>0.414 |
| **PH_V4** | **KE**<br>**PE** | 0.736<br>0.641 | 0.646<br>0.504 | 0.779<br>0.793 | 0.566<br>0.599 |
| **PH_V6** | **KE**<br>**PE** | 0.746<br>0.650 | 0.681<br>0.545 | 0.781<br>0.845 | 0.662<br>0.508 |
| **PH_final** | **KE**<br>**PE** | 0.688<br>0.668 | 0.768<br>0.739 | 0.721<br>0.790 | 0.778<br>0.771 |
| **FF** | **KE**<br>**PE** | 0.656<br>0.650 | 0.780<br>0.743 | -<br>- | 0.745<br>0.639 |
| **RL** | **KE**<br>**PE** | -<br>- | 0.436<br>0.226 | -<br>- | 0.377<br>0.329 |

**Table S3** Genomic correlation between 2017 and 2018 in each environment for trait EV_V3 for KE (blue numbers) and PE (red numbers). The blue and red bold numbers with stars indicate which proportion of interactions in bivariate sERRBLUP maximized the predictive ability in each environment for KE and PE, respectively.

| Bivariate Models | ROG | GOL | TOM |
|---|---|---|---|
| **GBLUP** | 0.864 / 0.442 | 0.984 / 0.768 | 0.913 / 0.642 |
| **sERRBLUP top 10%** | 0.869 / 0.643 | 0.804 / **0.740*** | **0.892*** / **0.704*** |
| **sERRBLUP top 5%** | 0.872 / 0.976 | **0.759*** / 0.725 | 0.877 / 0.686 |
| **sERRBLUP top 1%** | 0.844 / 0.947 | 0.760 / 0.824 | 0.896 / 0.765 |
| **sERRBLUP top 0.1%** | **0.884*** / **0.953*** | 0.812 / 0.931 | 0.920 / 0.997 |
| **sERRBLUP top 0.01%** | 0.829 / 0.968 | 0.781 / 0.973 | 0.993 / 0.938 |
| **sERRBLUP top 0.001%** | 0.857 / 0.940 | 0.892 / 0.962 | 0.962 / 0.908 |

**Table S4** Genomic correlation between 2017 and 2018 in each environment for trait EV_V4 for KE (blue numbers) and PE (red numbers). The blue and red bold numbers with stars indicate which proportion of interactions in bivariate sERRBLUP maximized the predictive ability in each environment for KE and PE, respectively.

| Bivariate Models | EIN | ROG | GOL | TOM |
|---|---|---|---|---|
| GBLUP | 0.843 / 0.525 | 0.897 / 0.592 | 0.940 / 0.998 | 0.960 / 0.835 |
| sERRBLUP top 10% | **0.810*** / 0.713 | **0.863*** / 0.618 | 0.872 / 0.815 | 0.972 / **0.744*** |
| sERRBLUP top 5% | 0.807 / 0.732 | 0.861 / 0.632 | 0.830 / **0.782*** | 0.997 / 0.751 |
| sERRBLUP top 1% | 0.865 / 0.769 | 0.889 / **0.705*** | **0.791*** / 0.824 | 0.941 / 0.733 |
| sERRBLUP top 0.1% | 0.966 / 0.888 | 0.936 / 0.766 | 0.874 / 0.946 | 0.934 / 0.784 |
| sERRBLUP top 0.01% | 0.979 / 0.908 | 0.947 / 0.780 | 0.883 / 0.999 | 0.979 / 0.806 |
| sERRBLUP top 0.001% | 0.974 / **0.898*** | 0.966 / 0.758 | 0.861 / 0.932 | **0.999*** / 0.834 |

**Table S5** Genomic correlation between 2017 and 2018 in each environment for trait EV_V6 for KE (blue numbers) and PE (red numbers). The blue and red bold numbers with stars indicate which proportion of interactions in bivariate sERRBLUP maximized the predictive ability in each environment for KE and PE, respectively.

| Bivariate Models | EIN | ROG | GOL | TOM |
|---|---|---|---|---|
| GBLUP | 0.768 / 0.712 | 0.989 / 0.965 | 0.900 / 0.940 | 0.703 / 0.764 |
| sERRBLUP top 10% | 0.817 / **0.646*** | 0.922 / **0.715*** | 0.895 / **0.886*** | **0.458*** / **0.594*** |
| sERRBLUP top 5% | **0.809*** / 0.635 | 0.900 / 0.736 | 0.861 / 0.883 | 0.412 / 0.566 |
| sERRBLUP top 1% | 0.809 / 0.690 | 0.888 / 0.818 | 0.823 / 0.898 | 0.409 / 0.542 |
| sERRBLUP top 0.1% | 0.890 / 0.842 | **0.936*** / 0.882 | **0.892*** / 0.942 | 0.489 / 0.544 |
| sERRBLUP top 0.01% | 0.991 / 0.840 | 0.954 / 0.908 | 0.914 / 0.916 | 0.496 / 0.530 |
| sERRBLUP top 0.001% | 0.909 / 0.899 | 0.969 / 0.968 | 0.856 / 0.932 | 0.567 / 0.573 |

**Table S6** Genomic correlation between 2017 and 2018 in each environment for trait PH_V6 for KE (blue numbers) and PE (red numbers). The blue and red bold numbers with stars indicate which proportion of interactions in bivariate sERRBLUP maximized the predictive ability in each environment for KE and PE, respectively.

| Bivariate Models | EIN | ROG | GOL | TOM |
|---|---|---|---|---|
| GBLUP | 0.942 / 0.880 | 0.952 / 1.000 | 0.937 / 0.952 | 0.994 / 0.758 |
| sERRBLUP top 10% | **0.932*** / **0.823*** | 0.909 / **0.727*** | 0.943 / **0.912*** | 0.949 / **0.608*** |
| sERRBLUP top 5% | 0.928 / 0.801 | 0.909 / 0.749 | 0.935 / 0.908 | **0.938*** / 0.568 |
| sERRBLUP top 1% | 0.910 / 0.823 | 0.951 / 0.826 | 0.887 / 0.934 | 0.936 / 0.501 |
| sERRBLUP top 0.1% | 0.934 / 0.915 | **0.991*** / 0.934 | **0.924*** / 0.957 | 0.955 / 0.964 |
| sERRBLUP top 0.01% | 0.902 / 0.874 | 0.959 / 0.963 | 0.969 / 0.948 | 0.968 / 0.526 |
| sERRBLUP top 0.001% | 0.918 / 0.848 | 0.984 / 0.955 | 0.919 / 0.941 | 0.993 / 0.713 |

**Table S7** Genomic correlation between 2017 and 2018 in each environment for trait PH_final for KE (blue numbers) and PE (red numbers). The blue and red bold numbers with stars indicate which proportion of interactions in bivariate sERRBLUP maximized the predictive ability in each environment for KE and PE, respectively.

| Bivariate Models | EIN | ROG | GOL | TOM |
|---|---|---|---|---|
| GBLUP | 0.831 / 0.851 | 0.984 / 0.935 | 0.958 / 0.943 | 0.985 / 0.981 |
| sERRBLUP top 10% | **0.822*** / **0.799*** | **0.986***/ 0.910 | **0.894*** / **0.871*** | **0.864*** / **0.971*** |
| sERRBLUP top 5% | 0.820 / 0.738 | 0.987 / 0.900 | 0.861 / 0.863 | 0.816 / 0.965 |
| sERRBLUP top 1% | 0.881 / 0.835 | 0.994 / 0.924 | 0.822 / 0.867 | 0.808 / 0.981 |
| sERRBLUP top 0.1% | 0.986 / 0.934 | 0.999 / 0.959 | 0.989 / 0.956 | 0.841 / 0.981 |
| sERRBLUP top 0.01% | 0.962 / 0.922 | 0.999 / 0.967 | 0.982 / 0.994 | 0.873 / 0.984 |
| sERRBLUP top 0.001% | 0.965 / 0.879 | 0.992 / **0.942*** | 0.988 / 0.998 | 0.935 / 0.987 |

**Table S8** Genomic correlation between 2017 and 2018 in each environment for trait FF for KE (blue numbers) and PE (red numbers). The blue and red bold numbers with stars indicate which proportion of interactions in bivariate sERRBLUP maximized the predictive ability in each environment for KE and PE, respectively.

| Bivariate Models | EIN | ROG | TOM |
|---|---|---|---|
| GBLUP | 0.888 / 0.951 | 1.000 / 0.979 | 0.995 / 0.970 |
| sERRBLUP top 10% | **0.871*** / 0.882 | 0.987 / 0.969 | **0.959*** / **0.955*** |
| sERRBLUP top 5% | 0.859 / **0.865*** | **0.984*** / 0.955 | 0.955 / 0.944 |
| sERRBLUP top 1% | 0.852 / 0.855 | 0.983 / **0.941*** | 0.995 / 0.993 |
| sERRBLUP top 0.1% | 0.854 / 0.861 | 1.000 / 0.955 | 0.987 / 0.981 |
| sERRBLUP top 0.01% | 0.847 / 0.909 | 0.997 / 0.970 | 0.957 / 0.977 |
| sERRBLUP top 0.001% | 0.814 / 0.898 | 0.992 / 0.961 | 0.992 / 0.984 |

**Table S9** Genomic correlation between 2017 and 2018 in each environment for trait RL for KE (blue numbers) and PE (red numbers). The blue and red bold numbers with stars indicate which proportion of interactions in bivariate sERRBLUP maximized the predictive ability in each environment for KE and PE, respectively.

| Bivariate Models | EIN | ROG |
|---|---|---|
| GBLUP | 0.662 / 0.818 | 0.751 / 0.404 |
| sERRBLUP top 10% | 0.672 / 0.604 | **0.753*** / **0.321*** |
| sERRBLUP top 5% | 0.644 / **0.520*** | 0.741 / 0.325 |
| sERRBLUP top 1% | 0.603 / 0.546 | 0.748 / 0.338 |
| sERRBLUP top 0.1% | **0.676*** / 0.719 | 0.887 / 0.412 |
| sERRBLUP top 0.01% | 0.727 / 0.572 | 0.973 / 0.380 |
| sERRBLUP top 0.001% | 0.712 / 0.531 | 0.983 / 0.352 |

**Table S10a** GBLUP predictive ability for prediction in 2018 with training the model either on 2018 data or the average phenotypic values of 2017 and 2018 in each environment for series of phenotypic traits in KE.

| Trait | Training set | EIN | ROG | GOL | TOM |
|---|---|---|---|---|---|
| **EV_V3** | 2018<br>2017 and 2018 average | NA<br>NA | 0.335<br>0.338 | 0.435<br>0.425 | 0.291<br>0.311 |
| **EV _V4** | 2018<br>2017 and 2018 average | 0.448<br>0.429 | 0.385<br>0.386 | 0.410<br>0.387 | 0.299<br>0.316 |
| **EV _V6** | 2018<br>2017 and 2018 average | 0.355<br>0.314 | 0.397<br>0.406 | 0.403<br>0.410 | 0.566<br>0.532 |
| **PH_V4** | 2018<br>2017 and 2018 average | 0.470<br>0.483 | 0.495<br>0.488 | 0.506<br>0.518 | 0.299<br>0.294 |
| **PH_V6** | 2018<br>2017 and 2018 average | 0.527<br>0.523 | 0.494<br>0.458 | 0.464<br>0.470 | 0.463<br>0.471 |
| **PH_final** | 2018<br>2017 and 2018 average | 0.554<br>0.542 | 0.597<br>0.631 | 0.513<br>0.526 | 0.656<br>0.635 |
| **FF** | 2018<br>2017 and 2018 average | 0.474<br>0.458 | 0.502<br>0.551 | 0.134<br>- | 0.496<br>0.506 |
| **RL** | 2018<br>2017 and 2018 average | 0.391<br>0.358 | 0.252<br>0.212 | -<br>- | -<br>- |

**Table S10b** GBLUP predictive ability for prediction in 2018 with training the model either on 2018 data or the average phenotypic values of 2017 and 2018 in each environment for series of phenotypic traits in PE.

| Trait | Training set | EIN | ROG | GOL | TOM |
|---|---|---|---|---|---|
| EV_V3 | 2018<br>2017 and 2018 average | NA<br>NA | 0.209<br>0.210 | 0.337<br>0.332 | 0.326<br>0.323 |
| EV _V4 | 2018<br>2017 and 2018 average | 0.185<br>0.205 | 0.261<br>0.269 | 0.625<br>0.646 | 0.363<br>0.371 |
| EV _V6 | 2018<br>2017 and 2018 average | 0.323<br>0.288 | 0.362<br>0.382 | 0.627<br>0.645 | 0.583<br>0.572 |
| PH_V4 | 2018<br>2017 and 2018 average | 0.464<br>0.480 | 0.510<br>0.396 | 0.641<br>0.661 | 0.437<br>0.439 |
| PH_V6 | 2018<br>2017 and 2018 average | 0.547<br>0.541 | 0.529<br>0.440 | 0.690<br>0.706 | 0.515<br>0.512 |
| PH_final | 2018<br>2017 and 2018 average | 0.620<br>0.547 | 0.537<br>0.370 | 0.682<br>0.699 | 0.580<br>0.572 |
| FF | 2018<br>2017 and 2018 average | 0.466<br>0.495 | 0.456<br>0.079 | 0.306<br>- | 0.401<br>0.363 |
| RL | 2018<br>2017 and 2018 average | 0.268<br>0.235 | 0.419<br>0.114 | -<br>- | -<br>- |

**Table S11** The traits heritabilities in 2017, 2018 and both years jointly in KE (blue numbers) and PE (red numbers).

| Traits | 2017 | 2018 | Both 2017 and 2018 |
|---|---|---|---|
| EV_V3 | 0.91 / 0.85 | 0.79 / 0.67 | 0.92 / 0.86 |
| EV_V4 | 0.90 / 0.82 | 0.83 / 0.71 | 0.91 / 0.84 |
| EV_V6 | 0.88 / 0.84 | 0.75 / 0.68 | 0.90 / 0.85 |
| PH_V4 | 0.89 / 0.82 | 0.81 / 0.72 | 0.92 / 0.87 |
| PH_V6 | 0.90 / 0.88 | 0.85 / 0.81 | 0.93 / 0.91 |
| PH_final | 0.91 / 0.93 | 0.83 / 0.85 | 0.94 / 0.94 |
| FF | 0.90 / 0.92 | 0.86 / 0.83 | 0.94 / 0.93 |
| RL | 0.78 / 0.55 | 0.49 / 0.29 | 0.80 / 0.59 |

**Table S12** The combinations which did not converged in bivariate models with the number of not converged folds in 5-fold cross validation with 5 replicates for all traits in KE (blue numbers) and PE (red numbers). The red and blue zeros in the parentheses represent the non-convergence of pre estimated variance components based on the full set in KE and PE, respectively.

| Traits | Predicted Environments | GBLUP | ERRBLUP | sERRBLUP Top 10% | sERRBLUP Top 5% | sERRBLUP Top 1% | sERRBLUP Top 0.1% | sERRBLUP Top 0.01% | sERRBLUP Top 0.001% |
|---|---|---|---|---|---|---|---|---|---|
| EV_V3 | ROG | 2 | 3 | 1 | 3 | 12(0) | 8(0) | 10(0) | 21(0) |
| | GOL | - | - | (0)25 | (0)25 | - | 7 | 4 / 14 | 1 / 10 |
| | TOM | - | - | 2 | 1 | - | 1 / 1 | 2 / 9 | (0)15 / 18(0) |
| | EIN | - | - | - | - | - | 5 / (0) | 6 | - |
| EV_V4 | ROG | 1 | 1 | - | - | 1 | - | 1 / 1 | 6 / 1 |
| | GOL | - | - | - | - | - | (0) | - | 8 |
| | TOM | (0)15 / 1 | 12 / 1 | 3 | - | 1 | (0) | - | 1 |
| | EIN | - | - | - | - | - | 1 | 1 | (0)/ 1 |
| EV_V6 | ROG | - | - | - | - | - | - | (0)11 / 2 | (0)25 |
| | GOL | - | - | (0)25 / 25(0) | 25 / 25(0) | (0)25 | (0)21 / 25(0) | 25(0) | - |
| | TOM | - | - | - | - | - | - | - | - |
| | EIN | - | - | - | 1 | - | - | 1 | - |
| PH_V4 | ROG | - | - | - | - | - | 4 | 4 | 2 |
| | GOL | - | - | (0)25 | (0)25 | (0)25 | (0)22 | - | - |
| | TOM | 1 / 2 | 1 / 1 | - | - | - | 2 | 2 | 1 |
| | EIN | - | - | - | - | - | - | - | - |
| PH_V6 | ROG | - | - | - | - | - | 4 | 2 | (0)4 |
| | GOL | 1 | 1 | 1 | 1 | - | - | 1 | (0)4 |
| | TOM | - | - | - | - | - | - | 1 | - |
| | EIN | - | - | - | - | 1 | 4 | (0)6 | 7(0) |
| PH_final | ROG | - | - | (0)25 / 3 | (0)25 / 25(0) | (0)25 / 25(0) | (0)25 / 25(0) | 2 / 25(0) | 13(0) |

| | | | | | | | | | |
|---|---|---|---|---|---|---|---|---|---|
| | **GOL** | - | - | - | - | 1 | - | - | 8 |
| | **TOM** | - | - | (0)24 / 1 | 2 | (0)20 / 1 | (0)24 / 8(0) | (0)1 / 12(0) | 6 |
| | **EIN** | - | - | 1 | - | 9 | 25(0) | 9 | 2 |
| **FF** | **ROG** | - | - | 1 / 25(0) | 25(0) | 1 / 25(0) | (0)24 / 25(0) | (0)11 / 25(0) | (0)19 |
| | **TOM** | 3 / 5 | (0)9 / 4 | 1 / 5 | 1 / 5 | 2 / 6 | 10 / 12(0) | (0)16 / 17(0) | 6 / 7 |
| **RL** | **EIN** | - | - | - | - | - | - | - | - |
| | **ROG** | - | - | - | - | 1 | 2 | 7 | 4 |

**Supplemental Figures**

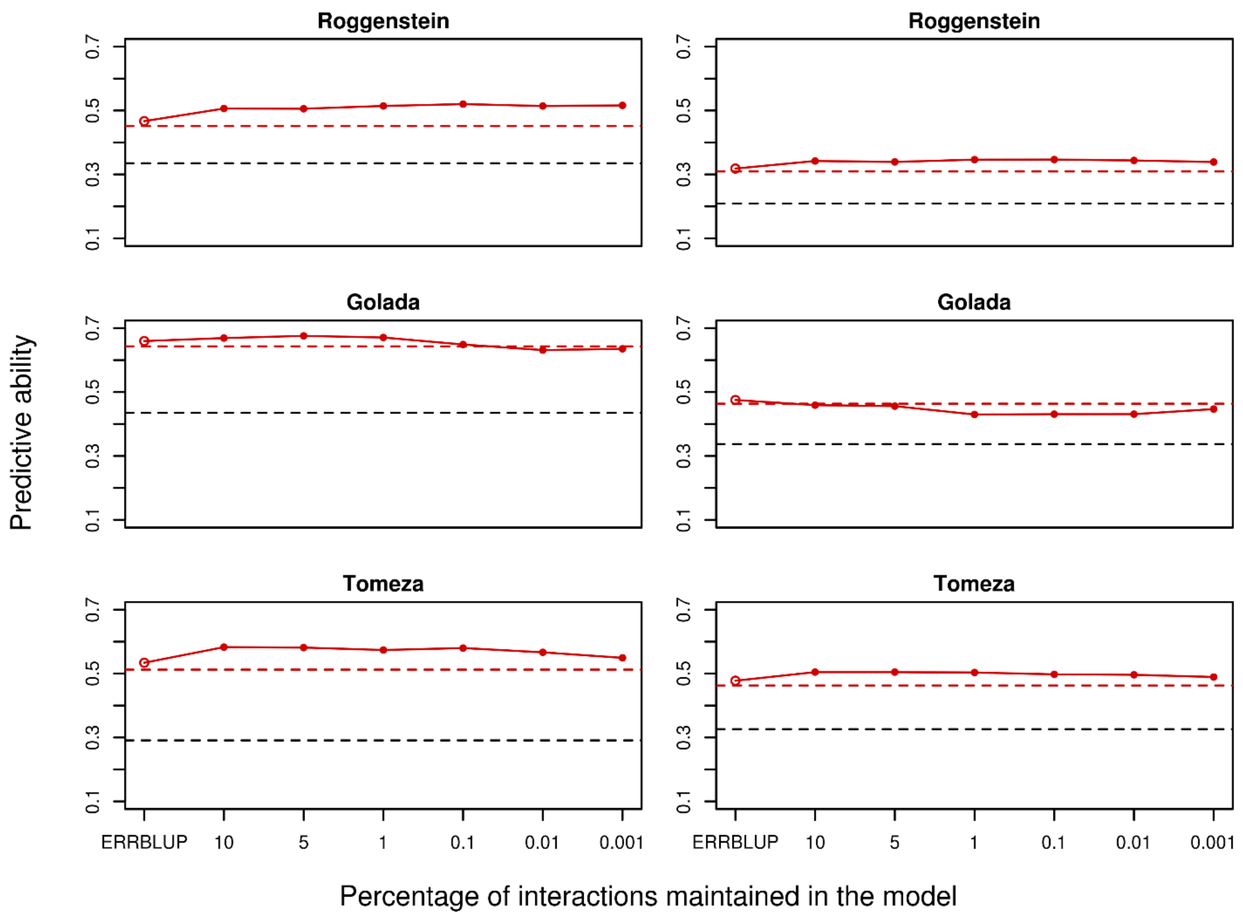

**Fig. S1** Predictive ability for univariate GBLUP within 2018 (black dashed horizontal line), bivariate GBLUP (red dashed horizontal line), bivariate ERRBLUP (red open circle) and bivariate sERRBLUP (red filled circles and red solid line) for trait EV_V3 in KE (left) and in PE (right).

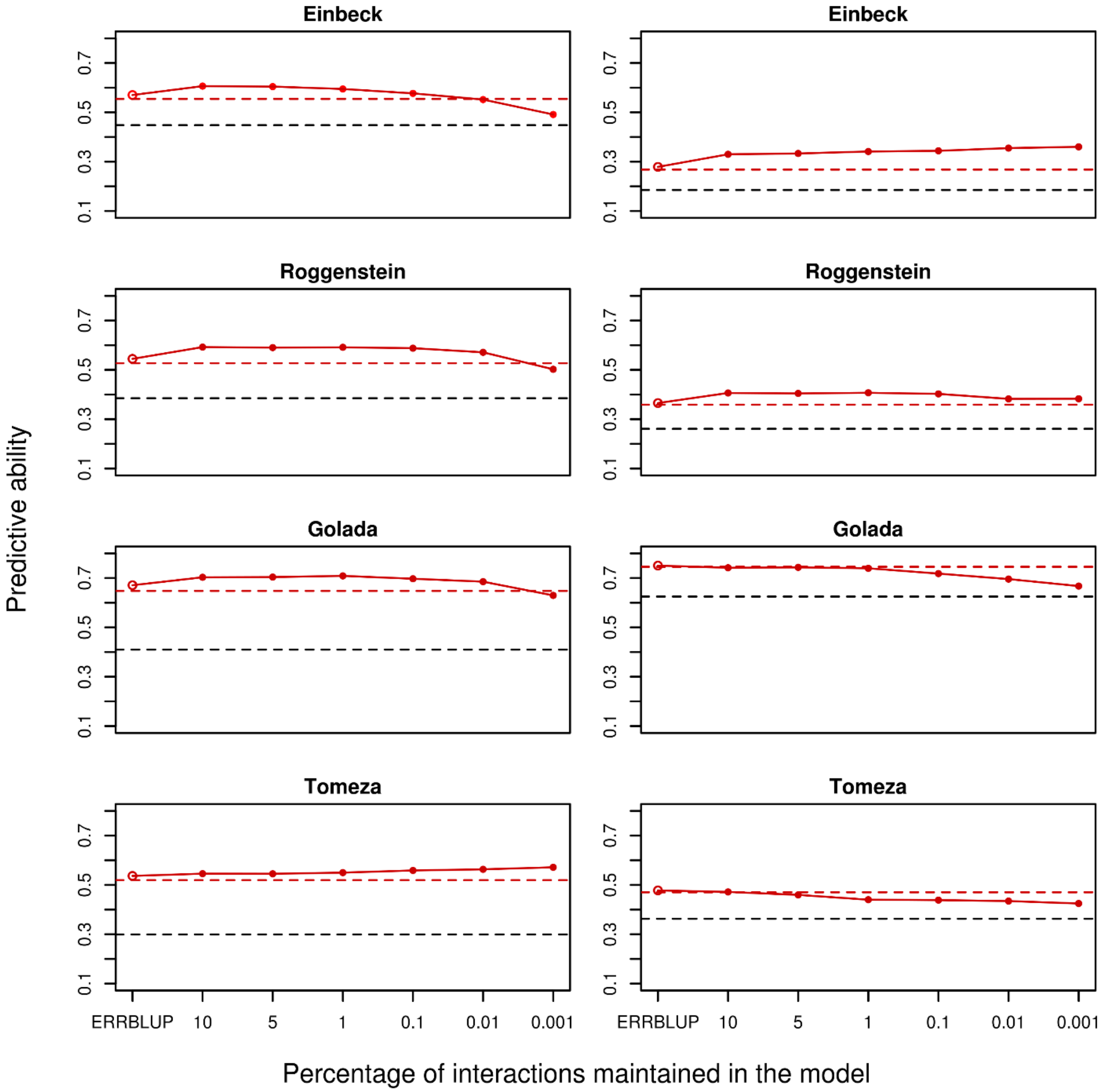

**Fig. S2** Predictive ability for univariate GBLUP within 2018 (black dashed horizontal line), bivariate GBLUP (red dashed horizontal line), bivariate ERRBLUP (red open circle) and bivariate sERRBLUP (red filled circles and red solid line) for trait EV_V4 in KE (left) and in PE (right).

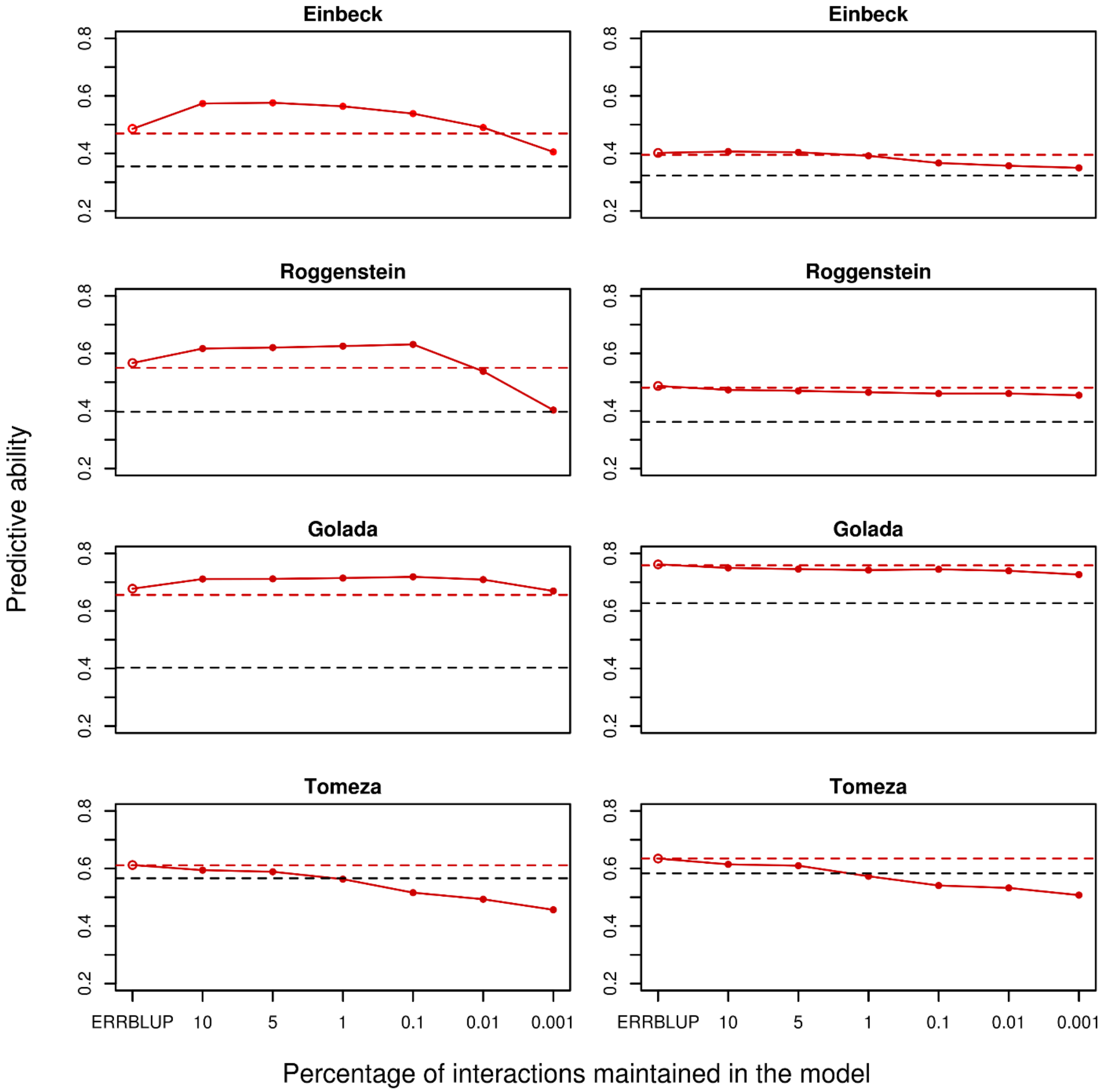

**Fig. S3** Predictive ability for univariate GBLUP within 2018 (black dashed horizontal line), bivariate GBLUP (red dashed horizontal line), bivariate ERRBLUP (red open circle) and bivariate sERRBLUP (red filled circles and red solid line) for trait EV_V6 in KE (left) and in PE (right).

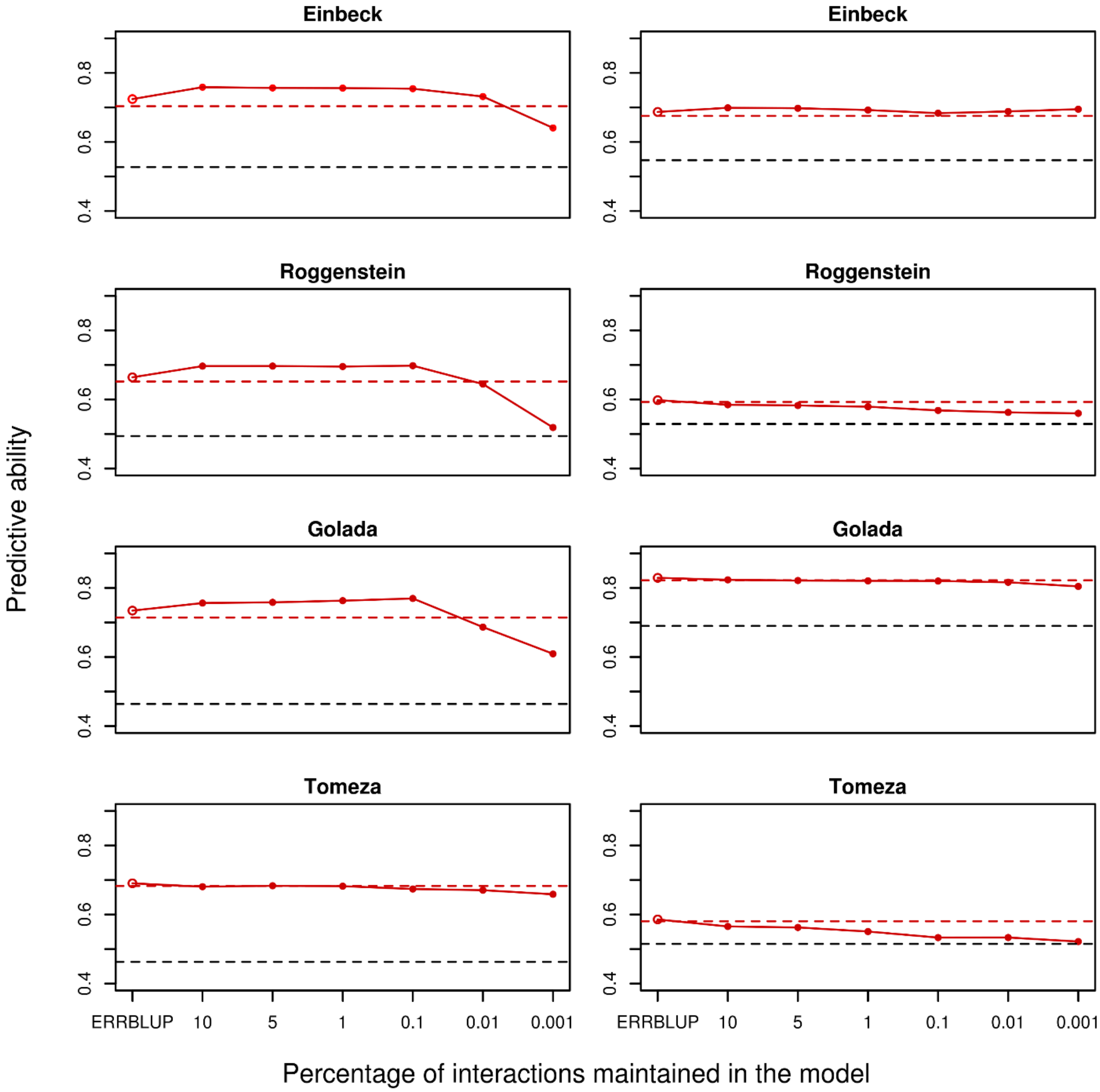

**Fig. S4** Predictive ability for univariate GBLUP within 2018 (black dashed horizontal line), bivariate GBLUP (red dashed horizontal line), bivariate ERRBLUP (red open circle) and bivariate sERRBLUP (red solid line) for trait PH-V6 in KE (left) and in PE (right).

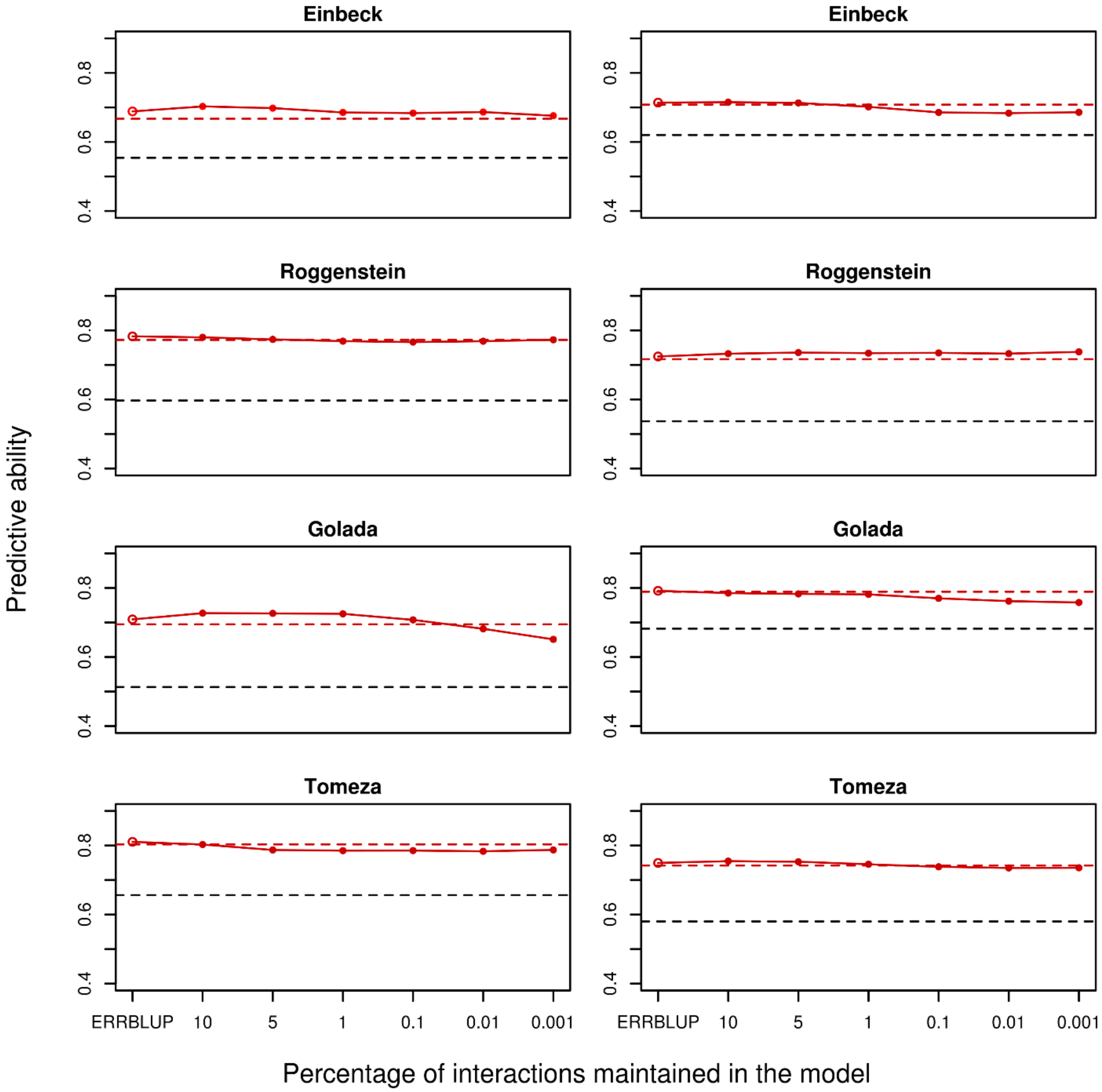

**Fig. S5** Predictive ability for univariate GBLUP within 2018 (black dashed horizontal line), bivariate GBLUP (red dashed horizontal line), bivariate ERRBLUP (red open circle) and bivariate sERRBLUP (red filled circles and red solid line) for trait PH-final in KE (left) and in PE (right).

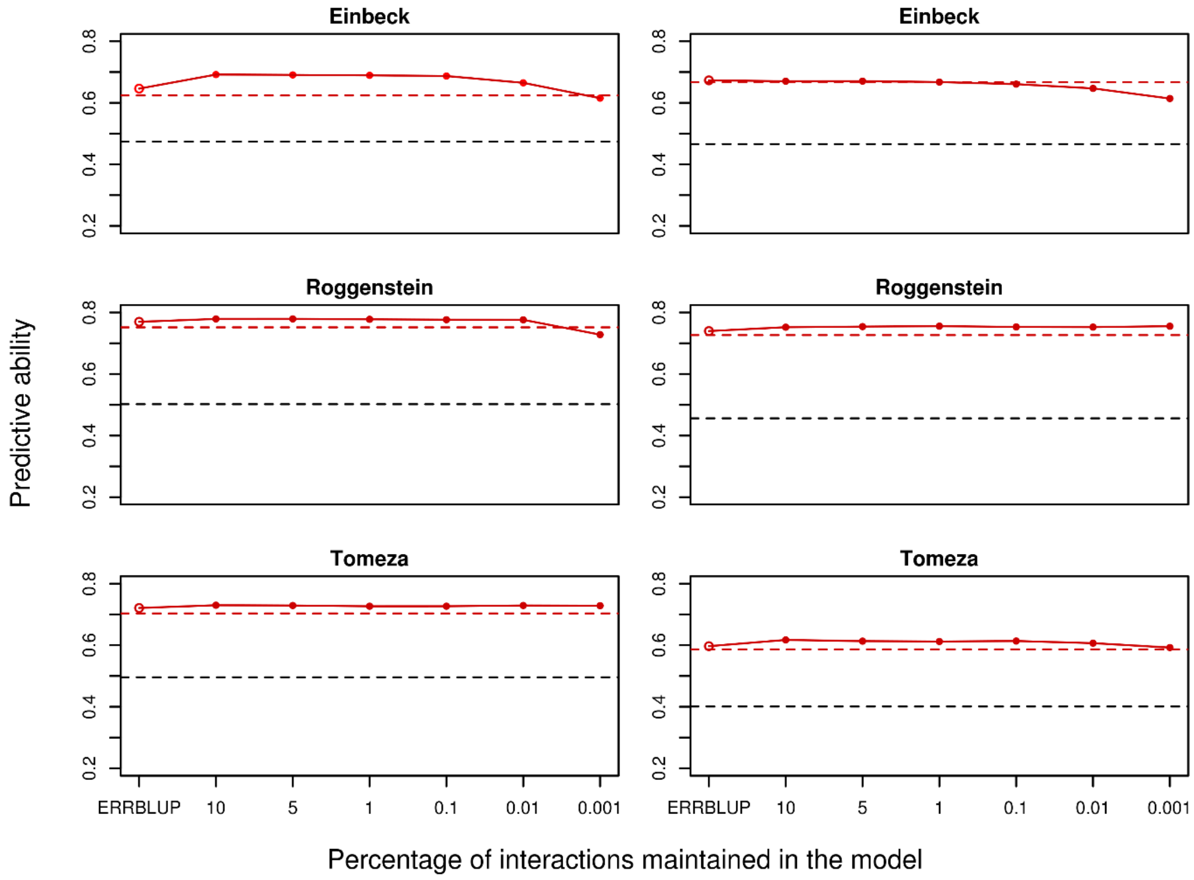

**Fig. S6** Predictive ability for univariate GBLUP within 2018 (black dashed horizontal line), bivariate GBLUP (red dashed horizontal line), bivariate ERRBLUP (red open circle) and bivariate sERRBLUP (red filled circles and red solid line) for trait FF in KE (left) and in PE (right).

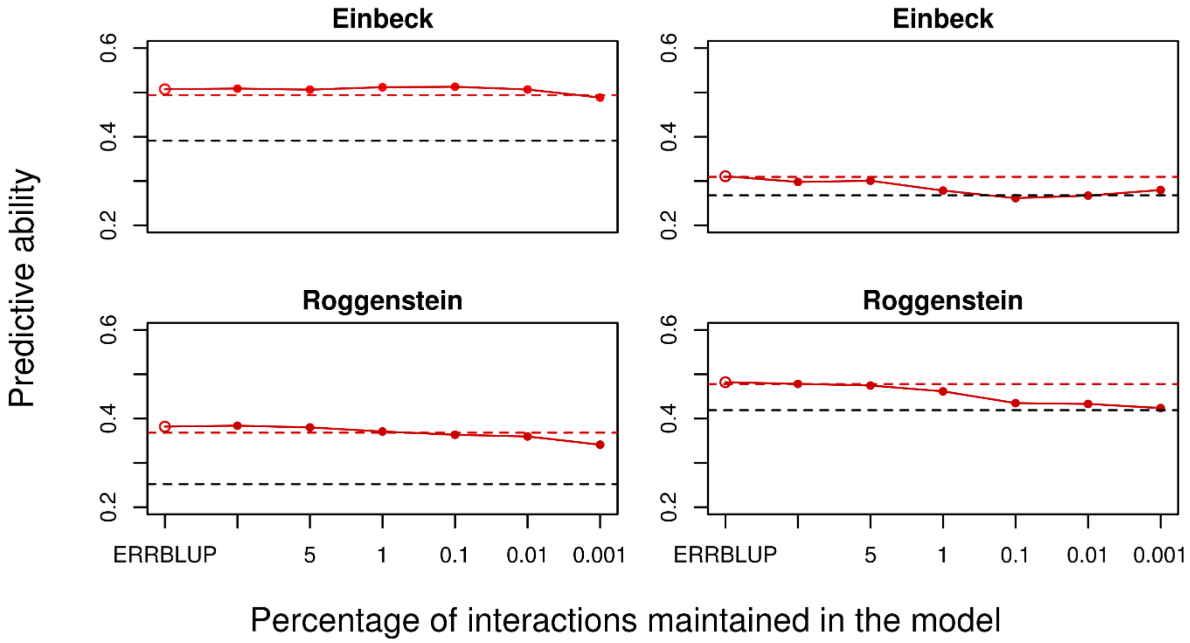

**Fig. S7** Predictive ability for univariate GBLUP within 2018 (black dashed horizontal line), bivariate GBLUP (red dashed horizontal line), bivariate ERRBLUP (red open circle) and bivariate sERRBLUP (red filled circles and red solid line) for trait RL in KE (left) and in PE (right).

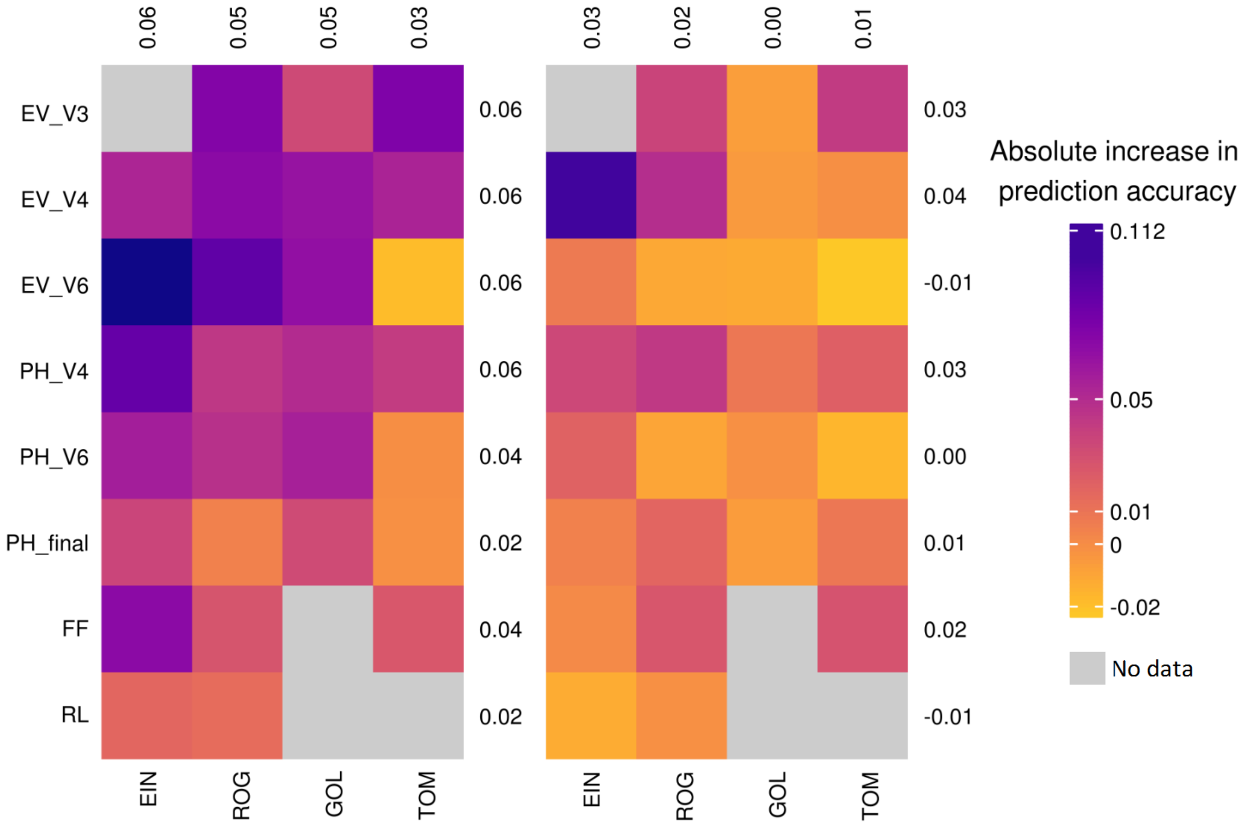

**Fig. S8** Absolute change in prediction accuracy from bivariate GBLUP to the maximum prediction accuracy of bivariate sERRBLUP in KE (left side plot) and in PE (right side plot). The average absolute change in prediction accuracy for each trait and environment is displayed in all rows and columns, respectively.

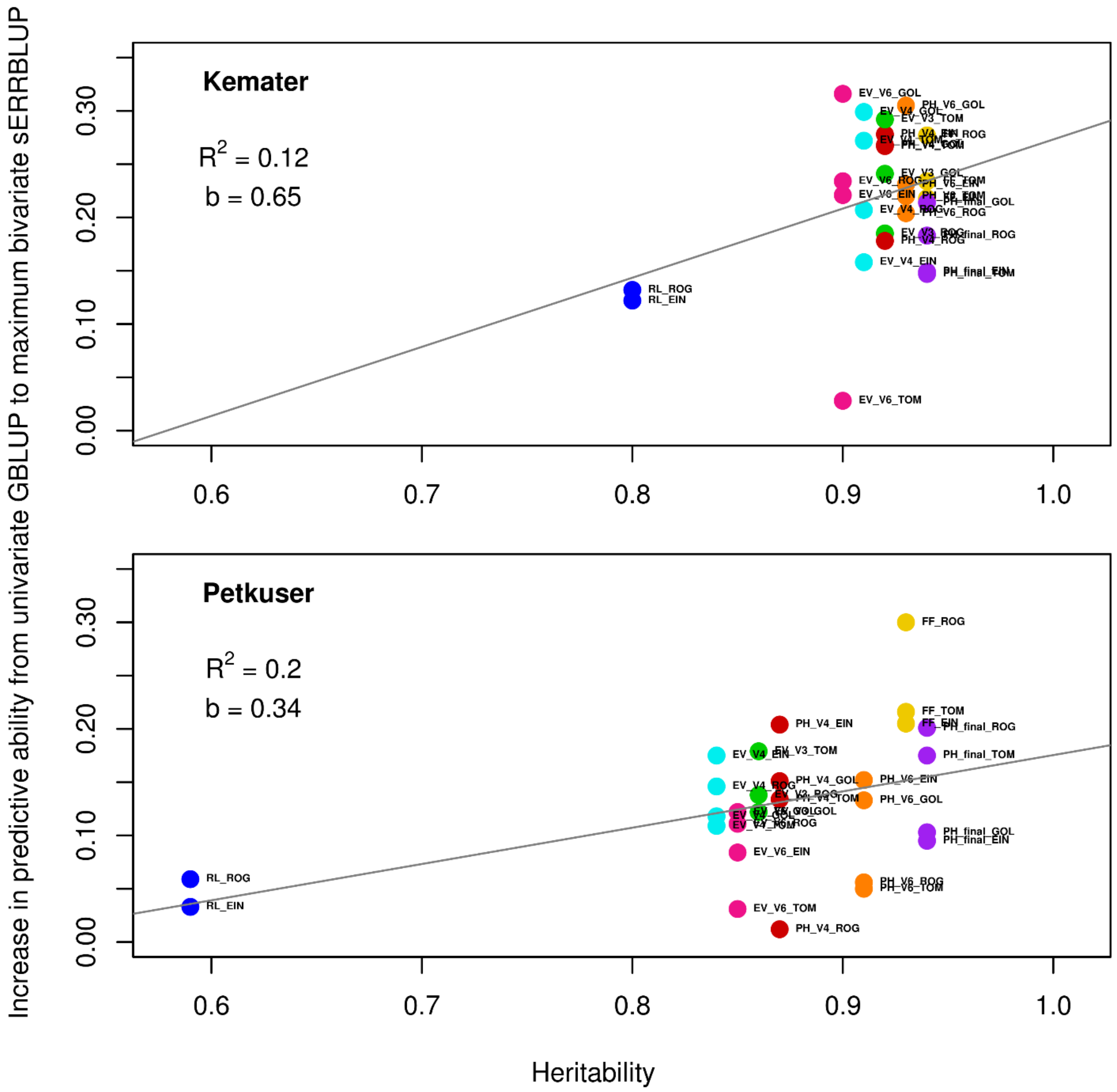

**Fig. S9** Regression of the absolute increase in predictive ability from univariate GBLUP to maximum bivariate sERRBLUP on the traits heritabilities calculated from both 2017 and 2018 in KE and in PE for all studied traits. In each panel, the overall linear regression line (gray solid line) with the regression coefficient ($\boldsymbol{b}$) and R-squared ($\boldsymbol{R^2}$) are shown.

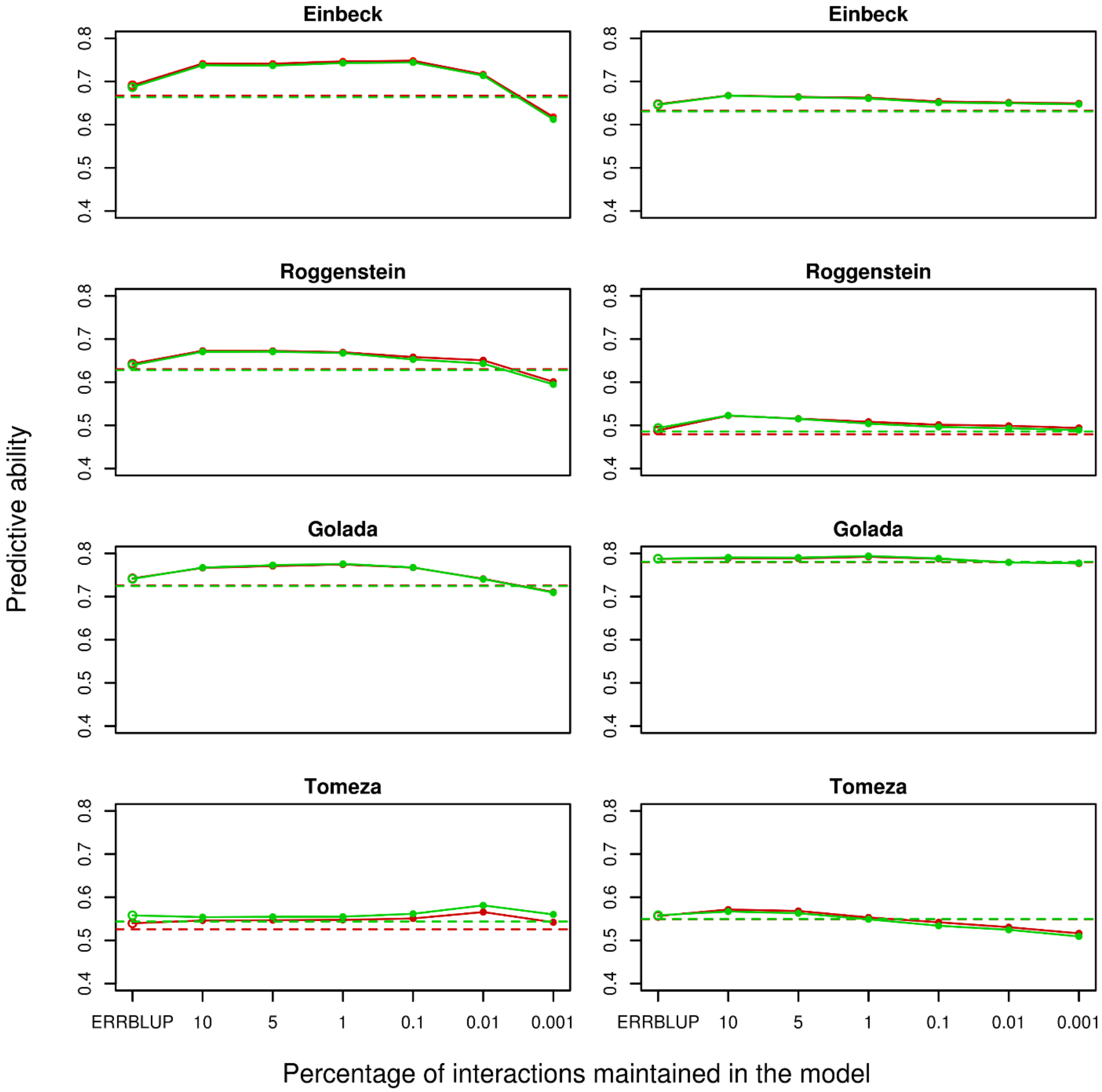

**Fig. S10** Comparison in predictive ability of bivariate GBLUP (dashed horizontal line), bivariate ERRBLUP (open circle) and bivariate sERRBLUP (filled circles and solid line) in 5-fold cross validation with 5 replicates (red circles and lines) and 10-fold cross validation with 10 replicates (green circles and lines) for trait PH-V4 in KE (left) and in PE (right).
